# RESPIRATION DEFECTS LIMIT SERINE SYNTHESIS REQUIRED FOR LUNG CANCER GROWTH AND SURVIVAL

**DOI:** 10.1101/2024.05.28.596339

**Authors:** Eduardo Cararo Lopes, Fuqian Shi, Akshada Sawant, Maria Ibrahim, Maria Gomez-Jenkins, Zhixian Hu, Pranav Manchiraju, Vrushank Bhatt, Wenping Wang, Christian S. Hinrichs, Douglas C. Wallace, Xiaoyang Su, Joshua D. Rabinowitz, Chang S. Chan, Jessie Yanxiang Guo, Shridar Ganesan, Edmund C. Lattime, Eileen White

## Abstract

Mitochondrial function is important for both energetic and anabolic metabolism. Pathogenic mitochondrial DNA (mtDNA) mutations directly impact these functions, resulting in the detrimental consequences seen in human mitochondrial diseases. The role of pathogenic mtDNA mutations in human cancers is less clear; while pathogenic mtDNA mutations are observed in some cancer types, they are almost absent in others. We report here that the proofreading mutant DNA polymerase gamma (*PolG*^*D256A*^) induced a high mtDNA mutation burden in non-small-cell lung cancer (NSCLC), and promoted the accumulation of defective mitochondria, which is responsible for decreased tumor cell proliferation and viability and increased cancer survival. In NSCLC cells, pathogenic mtDNA mutations increased glycolysis and caused dependence on glucose. The glucose dependency sustained mitochondrial energetics but at the cost of a decreased NAD+/NADH ratio that inhibited *de novo* serine synthesis. Insufficient serine synthesis, in turn, impaired the downstream synthesis of GSH and nucleotides, leading to impaired tumor growth that increased cancer survival. Unlike tumors with intact mitochondrial function, NSCLC with pathogenic mtDNA mutations were sensitive to dietary serine and glycine deprivation. Thus, mitochondrial function in NSCLC is required specifically to sustain sufficient serine synthesis for nucleotide production and redox homeostasis to support tumor growth, explaining why these cancers preserve functional mtDNA.

**In brief:** High mtDNA mutation burden in non-small-cell lung cancer (NSCLC) leads to the accumulation of respiration-defective mitochondria and dependency on glucose and glycolytic metabolism. Defective respiratory metabolism causes a massive accumulation of cytosolic nicotinamide adenine dinucleotide + hydrogen (NADH), which impedes serine synthesis and, thereby, glutathione (GSH) and nucleotide synthesis, leading to impaired tumor growth and increased survival.

**Highlights:**

- Proofreading mutations in Polymerase gamma led to a high burden of mitochondrial DNA mutations, promoting the accumulation of mitochondria with respiratory defects in NSCLC.
- Defective respiration led to reduced proliferation and viability of NSCLC cells increasing survival to cancer.
- Defective respiration caused glucose dependency to fuel elevated glycolysis.
- Altered glucose metabolism is associated with high NADH that limits serine synthesis, leading to impaired GSH and nucleotide production.
- Mitochondrial respiration defects sensitize NSCLC to dietary serine/glycine starvation, further increasing survival.

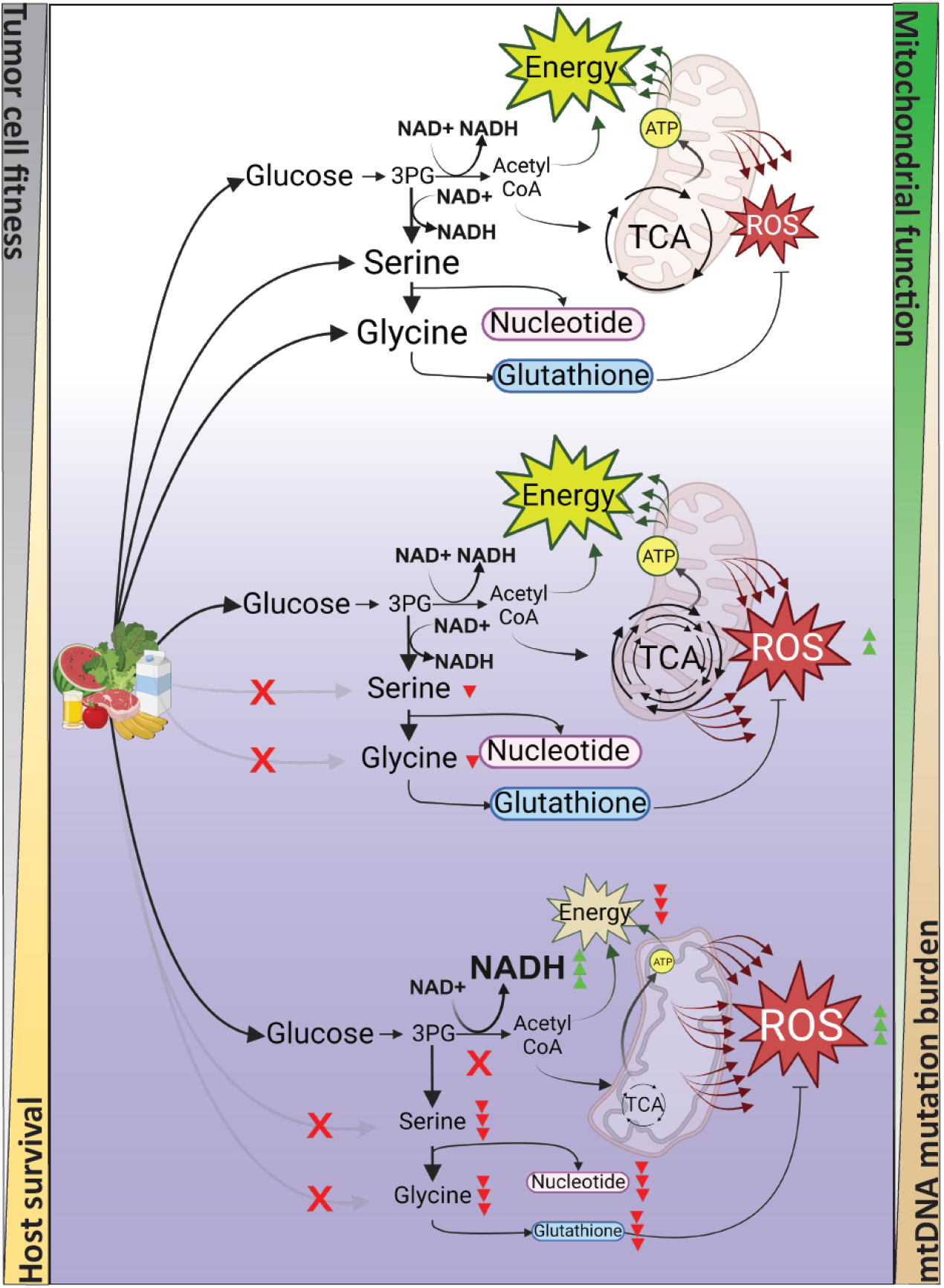

## INTRODUCTION

The balance between energetic (the breakdown of macromolecules to generate ATP) and anabolic (the production of macromolecules from smaller ones using energy) metabolism is a *sine quo-non* condition for tumor cell fitness^1^. While various cancer cells employ diverse strategies to achieve this balance, mitochondrial function plays a critical role in this equilibrium, particularly in certain cancers^2–4^. Inducing metabolic stress by limiting the availability or blocking the synthesis of specific nutrients in tumor cells can be an effective strategy for improving cancer therapies^5–9^. Therefore, understanding the metabolic mechanisms that support mitochondrial function and nutrient utilization for maintaining energy and anabolic balance is crucial. This insight will help identify specific targets to reduce tumor cell fitness without causing host toxicity.

Mitochondria serve as a prime example of this interplay between energetic and anabolic metabolism as they provide both energy and anabolic metabolites to sustain tumor cell growth and survival^10^. Nevertheless, only a limited number of cancer treatments target mitochondrial function, possibly due to concerns about potential adverse effects^11^. However, selectively inhibiting specific mitochondrial functions could sensitize tumor cells without compromising normal cellular function. Noteworthy examples of therapeutic strategies include studies involving the Complex I inhibitor metformin^12,13^, folate inhibitors^14^, nucleosides/nucleotides analogues^15^, and serine/glycine starvation.^16^ These strategies target key metabolic pathways: mitochondrial respiration, one-carbon (1C)-metabolism, nucleotide synthesis, and the serine synthesis pathway (SSP), pathways that play crucial roles in sustaining metabolic balance.^17–19^

One unique characteristic of mitochondria as organelles is that they harbor their own genetic material, the mitochondrial DNA (mtDNA). As mtDNA encodes essential proteins for the function of the electron transport chain (ETC), pathogenic mtDNA mutations can directly impact that important mitochondrial function^20–22^.

mtDNA mutations accumulate over time and are linked to mitochondrial dysfunction, which may play a role in mammalian aging^23,24^, and age represents the most significant risk factor for most cancers^25^. Consequently, it is reasonable to correlate an elevated mtDNA mutation burden with tumorigenicity^26^. However, pathogenic mtDNA mutations are found only in a few cancers, notably those of the colon, thyroid, and kidney.^4,27,28^ Despite the age-related accumulation of mtDNA mutations in the host, most cancers lack pathogenic mtDNA mutations^28^, presumably because these mutations may be deleterious to energy, anabolic, and redox homeostasis^29^. How specific cancers, including lung cancer, respond to an accumulation of mtDNA mutations and/or mitochondria dysfunction is unknown.

Lung cancer has a high mortality rate, affecting around 2.2 million people, leading to 1.8 million deaths worldwide annually^30^. Non-small-cell lung cancer (NSCLC) is the most common type of lung cancer^31^ and has been studied in the genetically engineered mouse model (GEMM) KP, (*Trp53*^*flox/flox*^; *KRas*^*LSL-G12D/+*^)^32^. The initiation of NSCLC in this mouse model can be conditionally induced by inhalation of adenovirus encoding Cre recombinase. This promotes the deletion of p53 (*Trp53*^Δ/Δ^) and activates the expression of the oncogenic *Kras* allele (*KRas*^G12D^). KP mimics the NSCLC naturally found in humans, including a relatively low level of mtDNA mutation burden^28,33^. Thus, to evaluate the impact of high mtDNA mutation burden in NSCLC, we generated a new GEMM, the PGKP (*PolG*^*D256A*/*D256A*^; *KRas*^*LSL-G12D/+*^; *Trp53*^*flox/flox*^) mouse model, by breeding KP with the PolG mutator mouse (*PolG*^*D256A*/*D256A*^) model^34^.

The PGKP animals can conditionally generate NSCLC as KP but contain a high mtDNA mutation burden due to a mutation in the polymerase gamma gene (*PolG*). PolG is a polymerase encoded by nuclear DNA (nDNA) but is responsible for replicating only mtDNA. The *PolG*^*D256A*^ mutation eliminates the proofreading exonuclease activity of this polymerase, dramatically reducing the fidelity of the polymerase and leading to the increased accumulation of mtDNA mutations over time. The PolG mutator mice have a whole-body *PolG*^*D256A*^ mutation causing accumulation of mtDNA mutations across all mouse tissues examined. The elevated mtDNA mutation burden contributes to various diseases linked to an aging phenotype. These diseases, such as severe erythroid dysplasia and megaloblastic anemia, result in a notably compromised body condition, ultimately reducing the lifespan of PolG mice to less than 18 months.^35–38^

To address the role of mtDNA mutation burden and mitochondrial function in NSCLC, we compared PGKP and KP mouse models following cancer induction. Introducing the *PolG* allele into KP lung cancers caused the accumulation of pathogenic mtDNA mutations and dysfunctional mitochondria, reduced tumor burden, and increased the lifespan of animals with NSCLC. This effect on tumor burden and survival was further increased when PGKP tumors were induced in older mice despite their poor body condition associated with the premature aging phenotype. Furthermore, mitochondrial dysfunction induced by mutated *PolG* led to defective respiration, prompting tumor cells to heavily rely on glucose and glycolysis. Moreover, respiratory defects resulted in the accumulation of NADH and a decreased NAD+/NADH ratio, leading to the suppression of the SSP and subsequent synthesis of glutathione (GSH) and nucleotides. Depleting dietary serine and glycine further repressed the growth of respiration-defective tumors while elevating dietary carbohydrates improved their growth. Finally, *in vitro* supplementation with GSH or nucleosides to compensate for serine/glycine starvation improved the growth of respiration-defective tumor cells, establishing why maintaining SSP is critical for tumor growth. Altogether, our findings demonstrate the essential role of mitochondrial function in supporting serine synthesis for nucleotides and redox homeostasis in NSCLC.

## RESULTS

### High mtDNA mutation burden increased lifespan in mice with NSCLC

The KP and PolG mouse models were crossed to generate PGKP mice with a constitutive germline mutation in PolG (PolGD256A/D256A), and Cre-inducible mutated *Kras* and floxed *Tp53* alleles (Figure 1A upper panel). Cohorts of young (2 months of age) and old (10 months of age) KP and PGKP animals were generated, NSCLC was induced, and the extent of mtDNA mutation burden in both tumor and normal lung tissue was examined. mtDNA sequencing of normal lung tissue from PGKP and KP pups (one day old) was performed to ascertain the initial mtDNA mutation burden (Figure 1A lower panel). Normal lung tissue and tumors were harvested from old and young mice 15 weeks post-NSCLC initiation (Figures 1A and S1A). Over time, PGKP animals accumulated greater numbers of single nucleotide variants (SNVs) and insertions/deletions (indels) in mtDNA in both normal lung tissue and NSCLC tumors in comparison to those in KP animals (Figure 1B). This accumulation was consistently higher in NSCLC tumors than in normal lung tissue of PGKP mice and was much higher than in KP tumors and normal lungs at all three-time points. Moreover, cytosine and guanosine misincorporation into mtDNA increased over time in PGKP tissues (Figure S1B), suggesting exposure to ROS or imbalanced nucleotide pools.^50^ In addition, most mtDNA genes in PGKP tumors accumulated mutations (Figure S1C), generating a similar mutational profile between young and old PGKP mice (Figure S1D).

**Figure 1.**
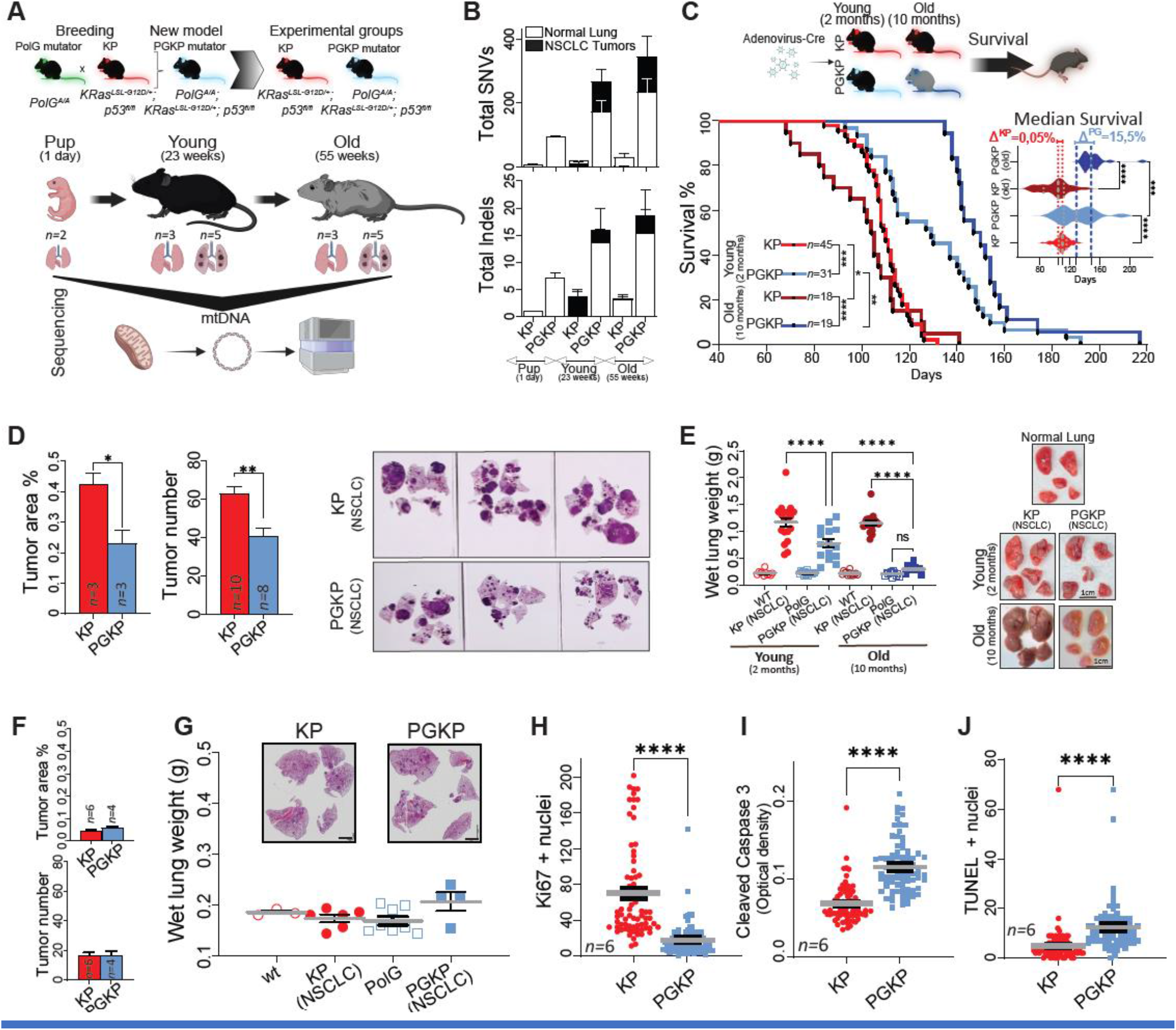
High mtDNA mutation burden limits NSCLC growth and malignancy without affecting tumor initiation. **(A)** At the top is a diagram illustrating the generation of the NSCLC conditional model with a high mtDNA mutation burden, PGKP. Below is a representation of mtDNA sequencing for KP and PGKP animals at various ages (*n* indicated per animal genotyping). **(B)** Shows the accumulation of mtDNA mutations, depicted by single nucleotide variants (SNVs) and Indels, in tissues and conditions as presented in panel A. **(C)** Survival curve of KP and PGKP, young (NSCLC initiation at two months of age) and old (NSCLC initiation at ten months of age) mice bearing NSCLC (check Figure S1A showing adeno-cre infection for NSCLC initiation). The median survival is illustrated to the right of the survival curve. **(D)** In young animals, tumor burden is indicated by tumor area (scanned slides) and number, 15 weeks after cancer initiation. H&E shows lung lobes affected by NSCLC in KP and PGKP animals during the same period. **(E)** Wet lung weight after 15 weeks of NSCLC initiation, with accompanying images illustrating normal lungs and NSCLC conditions. **(F-G)** Tumor area and number after six weeks of cancer initiation and wet lung weight in the same period. **(H-J)** IHC for ki67, cleaved caspase 3, and TUNEL assay were conducted on PGKP and KP tumors 15 weeks post NSCLC initiation (see Figure S2 for representative images). Data is shown as ± SEM. Two-tailed t-tests determined *P* values for D, F, H-J, while one-way ANOVA with Tukey correction was used for figures C, E, and G. Log-rank (Mantel-Cox) calculated P values for the survival curve. Significance levels are denoted as *≤0.05, **≤0.01, ***≤0.001, and ****≤0.0001.

The mutation frequency of mtDNA-encoded ETC genes was similar between young and old PGKP tumors (Figure S1E). These mutations might directly impact the activity of the ETC. To investigate this possibility, we used the sorting intolerant from tolerant mutations (SIFT) tool, which predicts whether an amino acid substitution affects the protein function^51^. In young and old animals, the predicted tolerated and deleterious mtDNA mutations were increased in PGKP compared with KP tumors (Figure S1G), suggesting a possible alteration of mitochondrial function.

To evaluate if the accumulation of mtDNA mutations influenced the survival of KP and PGKP models under cancer conditions, we initiated NSCLC in young animals. PGKP mice demonstrated a longer survival than KP mice (Figure 1C). This increased survival was attributed to reduced tumor burden, as evidenced by the reduction in tumor area and number by histology, hence reducing wet lung weight 15 weeks post NSCLC induction (Figures 1D-E).

As previously shown, the *PolG*^*D256A*^ mutation in PGKP mice leads to the accumulation of mtDNA mutations over time (Figure 1B)^34^. Therefore, we explored whether inducing NSCLC in old animals, which would be expected to have a greater mtDNA mutation burden at the time of cancer induction, would significantly impact their survival. To examine this possibility, cancer was initiated in old KP and PGKP mice. The survival in old PGKP with NSCLC was further increased compared to young PGKP mice, with the median survival being 15.5% higher than that of young PGKP mice with NSCLC (Figure 1C). The increased survival of old PGKP mice with NSCLC was due to remarkably reduced tumor burden 15 weeks post NSCLC induction (Figure 1E).

Although the model generated is just a tool to evaluate whether the accumulation of mtDNA mutations can impact mitochondria function and, consequently, NSCLC, we also tested mice carrying a single, homoplastic pathogenic mtDNA mutation associated with optic atrophy and Leigh Syndrome found in humans. This *MT-ND6* gene mutation (*MT-ND6*^*P25L*^) reduces complex I activity,^52^ and was introduced into KP mice to form ND6KP mice. Consistent with results from PGKP animals, the ND6KP mice also showed increased survival (17.5%) compared to KP mice after NSCLC induction (Figure S1G). Altogether, the results suggest that pathogenic mtDNA mutations are deleterious to NSCLC.

### mtDNA mutations impair NSCLC progression but not initiation

We hypothesized that the tumor burden in PGKP mice was lower than in KP mice because tumor initiation could be compromised, generating fewer malignant lesions after NSCLC induction. To test this hypothesis, NSCLC was induced in KP and PGKP mice, and the lungs of the animals were collected at six rather than 15 weeks post-tumor induction to assess the frequency of tumor initiation. In contrast to tumor burden at 15 weeks, at six weeks, there was little difference in tumor number and area by histology, and wet lung weight was also similar in both groups (Figures 1F-G).

Since we did not observe a difference in tumor initiation, we investigated whether tumor progression was affected by performing immunohistochemistry (IHC) for the proliferation marker Ki67, which was reduced in PGKP compared to KP tumors (Figures 1H, S2). Additionally, we evaluated tumor cell viability using IHC for cleaved caspase-3 and TUNEL assays. Both methods revealed elevated labeling in PGKP tumors, indicating lower viability than KP tumors (Figures 1I-J, S2). These results showed that the accumulation of mtDNA mutations reduced tumorigenesis by decreasing tumor cell proliferation and viability, likely contributing to the increased survival of PGKP mice with NSCLC.

### The *PolG* mutator allele elevates mtDNA mutations without affecting nuclear DNA

We investigated whether increased mtDNA mutation burden can indirectly promote mutations in nuclear DNA (nDNA), as this has been proposed as a means to promote cancer and aging.^23,53^ Since the tumors of PGKP animals presented with elevated levels of mtDNA mutations that increased with age (Figure 1B), we generated lung tumor-derived cell lines (TDCLs) from young and old PGKP and KP mice, and then we used these TDCLs to perform mtDNA genome and whole exome sequencing (WES) (Figure 2A). The levels of SNVs and indels in the mtDNA of PGKP TDCLs were substantially higher than those found in KP TDCLs, independent of whether the tumor cells were generated from young or old animals (Figures 2B-C). Moreover, despite the increased mtDNA mutation burden found in PGKP TDCLs, the levels of nDNA mutations are very similar between PGKP and KP TDCLs (Figures 2D-E). These findings suggest that impaired tumorigenesis of PGKP animals and possibly the aging process in PolG mutator animals results from alterations in mitochondrial function caused by pathogenic mutations in mtDNA and not due to alterations in nDNA.

**Figure 2.**
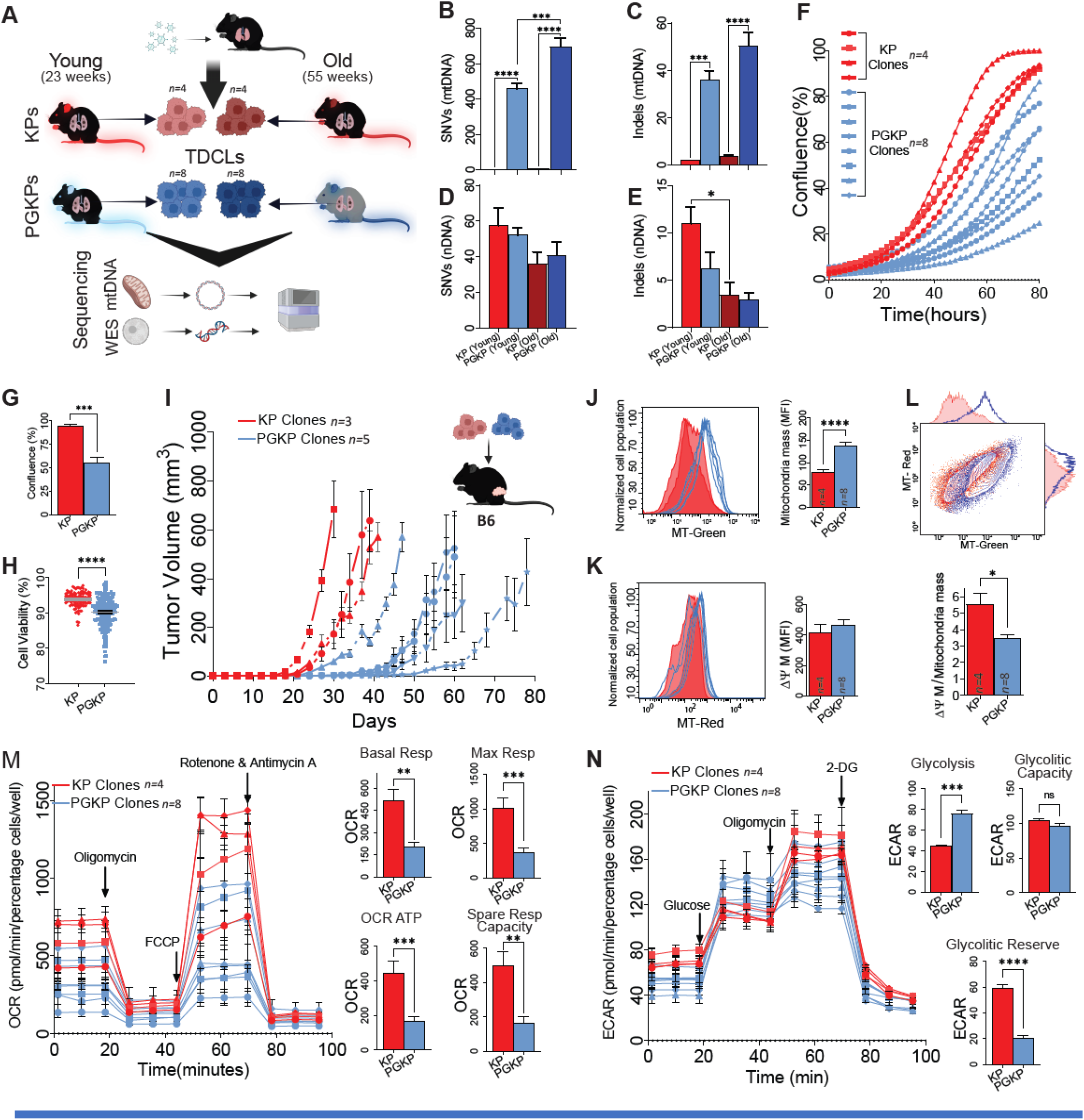
A high mtDNA mutation burden induces dysfunctional mitochondria and glycolytic addiction without impacting nuclear DNA mutations. **(A)** Schematic illustrating the generation of TDCLs from NSCLC tumors in KP, PGKP, young, and old animals. All TDCLs were independently derived from distinct tumors and subjected to mtDNA genome sequencing and whole exon sequencing. The figure displays the number of independent clones per group. **(B-C)** The mtDNA mutation burden is represented by SNVs and Indels of mtDNA in PGKP and KP TDCLs. **(D-E)** The nDNA mutation burden is represented by SNVs and Indels of mtDNA in PGKP and KP TDCLs. **(F)** Growth curve of KP and PGKP TDCLs (Incucyte) derived from young animals. **(G)** The confluence of TDCLs at the end of 80h. **(H)** The viability of the TDCLs was measured during 13 passages. Each point represents one viability test per clone. **(I)** Allograft tumor growth assay of KP and PGKP TDCLs in C57/B6 mice. **(J-L)** Flow cytometry assessing mitochondrial mass and mitochondrial membrane potential in TDCLs derived from KP and PGKP. **(I)** Oxygen consumption rate (OCR) and extracellular acidification rate (ECAR) were measured to assess mitochondrial respiration and glycolytic function in TDCLs. “*n*” denotes the number of independent clones utilized in the experiment. Carbonyl cyanide-4 (trifluoromethoxy) phenylhydrazone (FCCP) and 2-deoxy-glucose (2-DG) were employed in the study. Data is shown as ± SEM. Two-tailed t-tests determined *P* values for G, H, J, K, L, M, and N, while one-way ANOVA with Tukey correction was used for figures B, C, D, and E. Significance levels are denoted as *≤0.05, **≤0.01, ***≤0.001, and ****≤0.0001.

### High mtDNA mutation burden in NSCLC leads to the accumulation of dysfunctional mitochondria

To address whether the defective tumor growth seen in PGKP mice was tumor cell-intrinsic rather than a host effect of the *PolG* mutation in the germline, we evaluated the growth and survival of PGKP and KP TDCLs *in vitro*. PGKP TDCLs exhibited reduced proliferation and viability compared to that from KP (Figures 2F-H).

Furthermore, PGKP TDCLs demonstrated slower growth than KP TDCLs when implanted into syngenetic C57B/6 mice that lack the *PolG*^*D256A*^ mutator allele (Figure 2I). This indicates that PGKP NSCLC tumor cells inherently possess defects in cell proliferation and viability, replicating the same phenotype observed in induced PGKP tumors (Figures 1H-J, S2). It is noteworthy that the PGKP phenotype persisted irrespective of the experimental strategy employed, whether *in vitro* or *in vivo*, and whether conditional or allograft models were utilized.

We used flow cytometry to investigate the mitochondrial mass and membrane potential of NSCLC cells. The PGKP TDCLs had increased mitochondrial mass compared with those from KP (Figure 2J). However, their mitochondrial membrane potential was similar (Figure 2K), suggesting that PGKP tumor cells accumulate dysfunctional mitochondria (Figure 2L).

To directly evaluate mitochondrial function, we measured the oxygen consumption rate (OCR) in TDCLs (Figure 2M). As suggested by cytometry data (Figure 2L), mitochondrial function in PGKP tumor cells was compromised, presenting with a lower basal and maximum respiration expected to correspond to lower production of ATP by oxidative phosphorylation (OXPHOS). Moreover, the lack of spare respiratory capacity suggests permanent damage to the ETC of PGKP TDCLs.

Tumor cells with lower mitochondrial respiration usually rely on glycolysis to produce ATP and sustain their growth.^54,55^ We, therefore, measured the extracellular acidification rate (ECAR) of TDCLs (Figure 2N). As expected, the glycolytic activity was increased in PGKP compared to KP TDCLs. Interestingly, PGKP TDCLs had a much lower level of glycolytic reserve than KP TDCLs, indicating maximizing glycolysis in PGKP cells for their metabolic needs. These results demonstrate that the elevated mtDNA mutation burden in NSCLC cells results in the accumulation of dysfunctional mitochondria with respiration defects and enhanced glycolysis.

### PolG mice deplete circulating glucose, which contributes to defective tumor growth

Independent of the tumor cell-intrinsic growth and fitness defect caused by the *PolG* mutation, the physiological changes associated with mitochondrial dysfunction in PGKP mice may cause a systemic metabolic deficit that impairs tumor growth. To test this hypothesis, we assessed metabolic parameters in blood plasma and solid tissues (Figure 3A) in mice with NSCLC, comparing PGKP with KP in young and old mice. As controls for PGKP and KP, we utilized cancer-free animals, specifically PolG (*PolG*^*D256A/D256A*^; *Trp53*^*fl/fl*^) and *PolG* wild type (*Trp53*^*fl/fl*^), respectively. Both groups were also infected with adeno-Cre, but the control animals did not develop NSCLC due to the lack of *KRas*^*LSL-G12D/+*^. The lower blood glucose levels, post 15 weeks of NSCLC initiation, were observed in KP animals with NSCLC compared with control wild-type mice (Figure 3B). In PGKP and PolG animals, the alteration in blood glucose levels was more pronounced and had a clear age component, reinforcing that further accumulation of pathogenic mtDNA mutations can impact mutator mouse physiology and, consequently, NSCLC malignancy. It was reported that the PolG mutator mouse exhibits hypoglycemia in a starved condition.^56^ However, we observed that the blood glucose levels of PolG and PGKP, mainly in old animals, were within the range of hypoglycemia even in a fed state (Figure 3B).^57^

**Figure 3.**
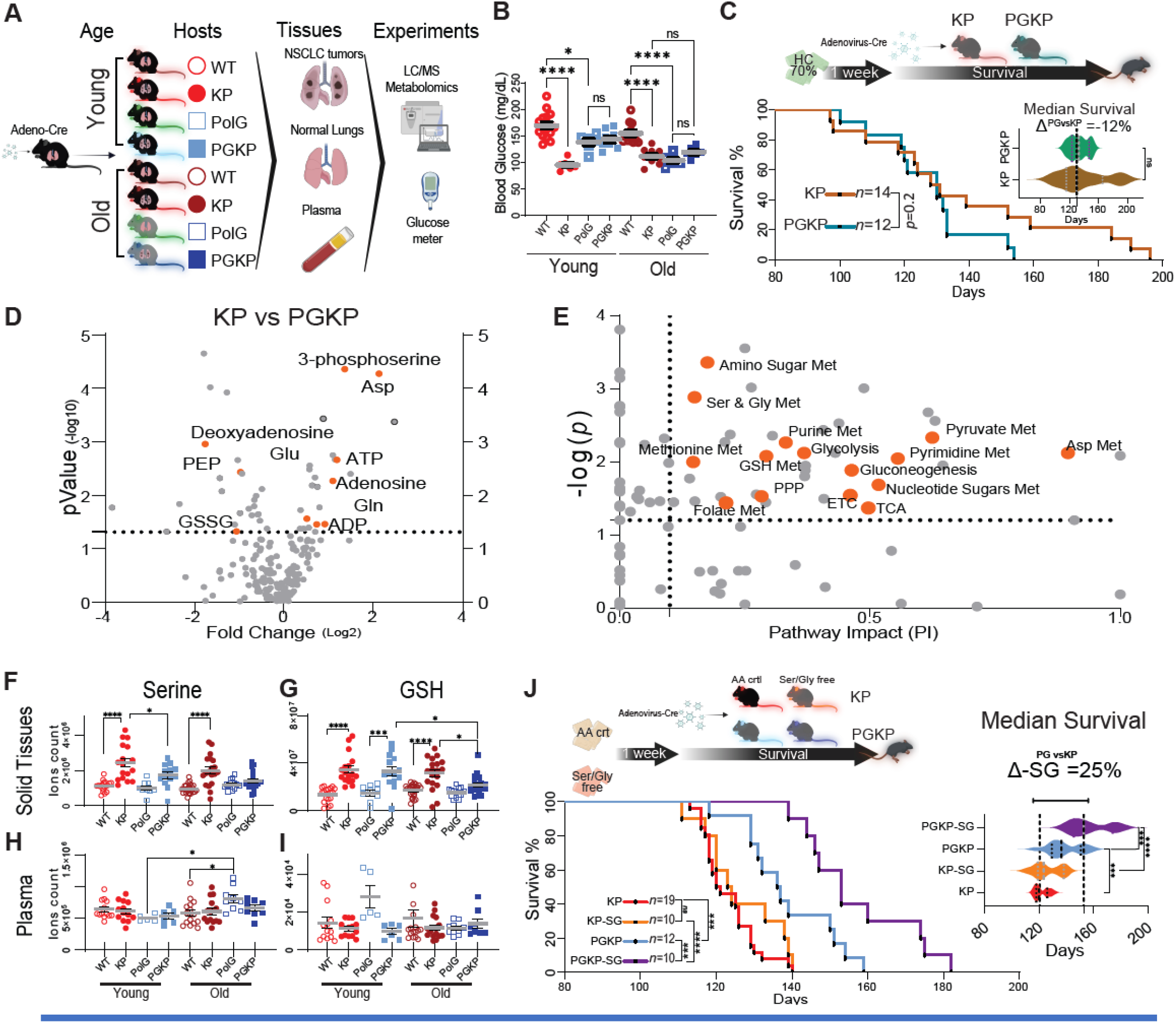
Tumor respiration defect sensitizes NSCLC to serine/glycine starvation: **(A)** Experimental design schematic: Old and young KP, WT, PGKP, and PolG animals were infected with adenovirus-Cre (KP and PGKP). After 15 weeks, euthanasia was performed, and plasma, NSCLC tumors (only KP and PGKP develop NSCLC), and normal lungs (from WT and PolG) were collected. The gathered material underwent pool-size metabolomics and glucose meter measurements. **(B)** Blood glucose levels. **(C)** Survival curve and median survival of KP and PGKP animals under a high carbohydrate diet (70% sucrose). **(D-E)** Representative volcano plot and pathway impact analysis for pool-size metabolomics comparing old KP and PGKP tumors. **(F-G)** Serine and GSH levels are shown for normal lungs (WT and PolG) and NSCLC tumors (PGKP and KP) at different ages. **(H-I)** Serine and GSH plasma levels in the same animals. **(J)** Survival curve and median survival of KP and PGKP animals under amino acid control (AA-crtl) and serine/glycine-free diet. Data is shown as ± SEM. Two-tailed t-tests determined *P* values for C (median survival), while one-way ANOVA with Tukey correction was used in B, C, F, G, H, I, and J. Log-rank (Mantel-Cox) calculated P values for survival curves C and J. Significance levels are denoted as *≤0.05, **≤0.01, ***≤0.001, and ****≤0.0001.

We hypothesized that the metabolic changes imposed by the accumulation of mtDNA mutations and defective mitochondria in PGKP animals could decrease glucose availability to tumors, thereby contributing to a reduction in tumor growth. Therefore, we tested if dietary manipulation of glucose would rescue tumor growth and survival defects caused by PolG by using a high carbohydrate (HC) diet composed of 70% sucrose (Figure 3C). We found that the HC diet eliminated the survival difference observed between PGKP and KP animals with NSCLC (Figure 1C), suggesting that systemic glucose limitation of highly glycolytic tumors and mouse tissues may contribute to the defective growth of NSCLC.

### Accumulation of defective mitochondria sensitizes NSCLC to dietary serine/glycine starvation

To further evaluate the underlying metabolic alterations impacting NSCLC tumor growth, we performed untargeted metabolomic profiling on tumors from PGKP and KP animals and their respective controls (Figure 3A). Remarkably, when compared to KP tumors, PGKP tumors exhibited significant alterations in 3-phosphoserine, an SSP precursor that uses glycolytic intermediates (Figure 3D). Furthermore, we observed reduced levels of aspartate in PGKP tumors, which serves as a biomarker of electron transport chain inhibition. This can be caused by the low ratio of NAD+/NADH, which promotes oxaloacetate reduction to malate, rendering oxaloacetate unavailable as a substrate for aspartate synthesis.^58^ Consequently, aspartate deficiency may pose a challenge to nucleotide synthesis. Indeed, the pathway impact analysis showed that the alterations in PGKP compared with KP tumors were concentrated in energetic (glycolysis and the TCA cycle) and anabolic (serine/glycine, aspartate metabolism, purine/pyrimidine, nucleotide sugar, glutathione (GSH), and 1C-metabolism) (Figure 3E).

Considering the alterations in anabolic metabolism identified in the pathway impact analysis and the diminished levels of 3-phosphoserine in PGKP tumors, we sought to identify the underlying metabolic mechanism and whether this contributed to defective tumor growth. Serine levels in NSCLC tumors from KP mice were higher than those in normal lung tissue from wild-type mice at any age (Figure 3F). However, the difference in serine levels in tumors from PGKP animals and normal lung tissue in PolG animals did not reach significance (Figure 3F). This suggests that an elevated level of serine in tumors is a distinctive feature of KP NSCLC that was absent from PGKP tumors. Similarly noteworthy were the changes in GSH, mirroring the trend observed in serine but with a more pronounced reduction in GSH levels in PGKP tumors from old mice compared to normal lungs from old PolG and old KP animals (Figure 3G). These variations were not evident in the plasma, suggesting that alterations in serine and GSH are characteristics of tumors (Figure 3H-I). These results suggested that because of limited substrate levels (e.g. glucose), PGKP tumor cells may ineffectively synthesize important downstream metabolites such as amino acids to sustain fitness. In turn, this could make PGKP tumors more sensitive to reduction in external key nutrients such as serine. To test this hypothesis, we fed KP and PGKP animals with a control amino acid diet (AA-crt) and a serine/glycine-free diet and monitored the survival of these mice after NSCLC initiation. The KP animals fed with these different diets had similar median survival (Figure 3J). However, PGKP animals with NSCLC had a 16% improved survival under serine/glycine starvation compared to PGKP animals on the AA-crtl diet and 25% compared with KPs in both diets (Figure 3J). Together, these results suggest that mitochondrial dysfunction in PGKP tumors leads to insufficient serine synthesis using glucose as substrate, creating an increased dependency on an exogenous supply of serine from the diet to support tumor growth.

### PGKP tumors produce less serine from glucose

Serine is a non-essential amino acid that can be synthesized from different sources, including glucose, and is also obtained from dietary sources (Figure 4A).^59^ As previously observed, the circulating glucose in the plasma and serine levels in tumors of PGKP animals were lower than those in KP animals. Therefore, we investigated whether there was a difference in how KP and PGKP tumors used glucose to make serine. Wild type, PolG, KP, and PGKP animals were pre-fed with an AA-crt or ser/gly-free diet for one week; tumor formation was then induced in the KP and PGKP animals, and the animals were continued on the same diets for an additional 12 weeks. We then performed *in vivo* isotope tracing by infusing [U-^13^C]-Glucose into these animals and measured the metabolic alterations in different tissues (Figure 4B).^60^

**Figure 4.**
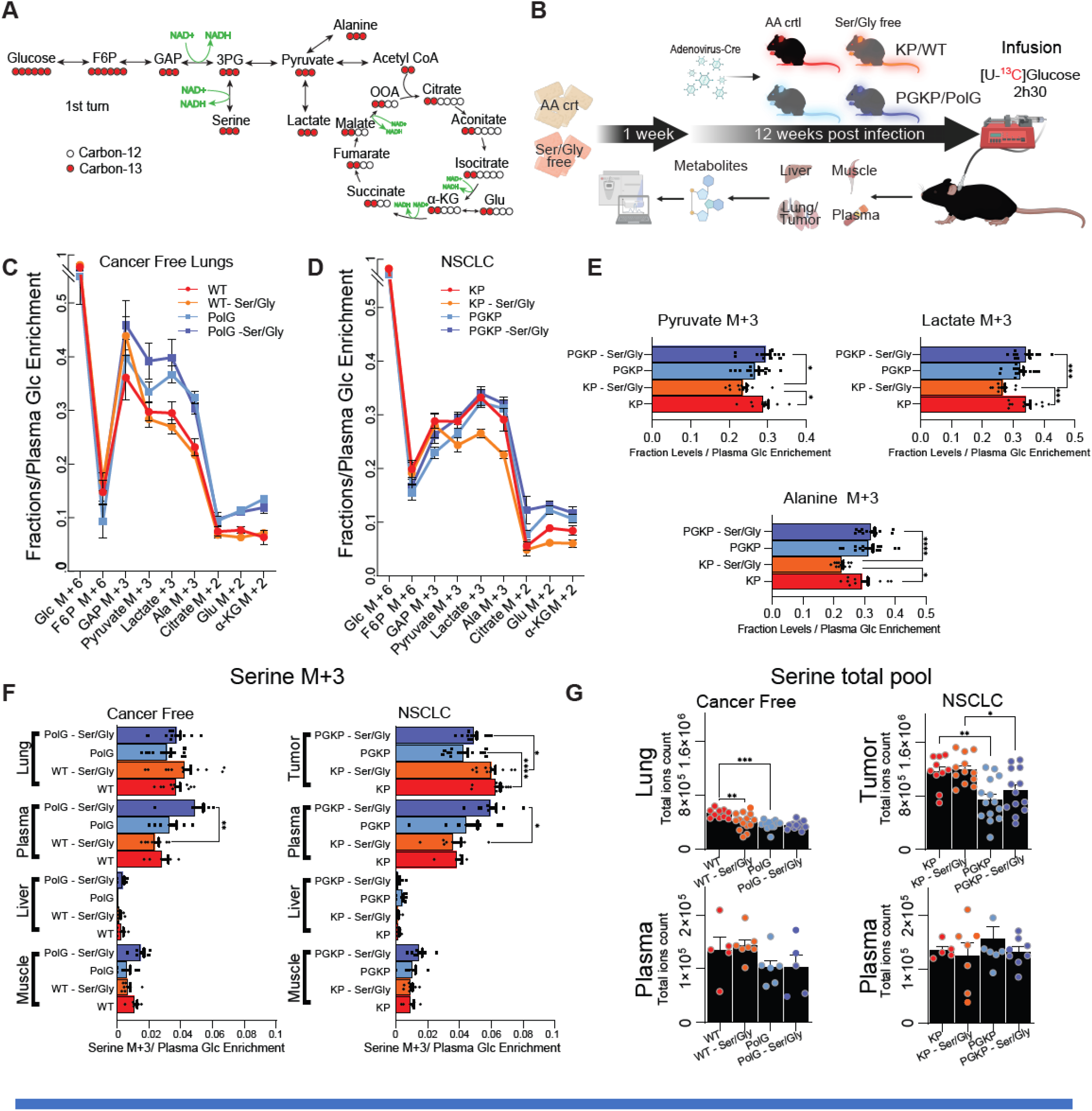
NSCLC rewires glycolytic metabolism to synthesize more serine from glucose: **(A)** Schematic depicting glycolysis→TCA with ^13^C distribution on pathway intermediates using [U-^13^C]-glucose. Intermediates include Fructose 6-phosphate (F6P), glyceraldehyde 3-phosphate (GAP), 3-phosphoglycerate (3PG). **(B)** Schematic experimental design for in vivo isotope tracing with [U-^13^C]-glucose infusion. KP, PGKP, WT, and PolG animals were infected with adenovirus-Cre and maintained on AA-crt or serine/glycine-free diets for 12 weeks. After fasting for six hours, animals underwent 2h30 of [U-^13^C]-glucose infusion and were then euthanized. Muscle, liver, plasma, and tumors (two independent tumors per animal) or lungs (two lobes per animal) were snap-frozen. The collected samples were subjected to pool-size metabolomics (n=5-7 animals per group), and tissue metabolite enrichment or isotopologue was normalized by the plasma glucose enrichment of the corresponding animal. **(C-D)** Dilution of ^13^C across the first cycle of glycolysis→TCA intermediates in normal (cancer-free) lungs and NSCLC tissues of mice infused with [U-^13^C]-glucose. **(E)** Statistical analysis of the main altered metabolites enrichment presented in D. **(F)** Serine M+3 fractions (first turn of glycolysis→SSP). **(G)** Total pool size levels of serine in lung, tumor, and their matched plasma. Data is shown as ± SEM. For all graphs the *P* values were determined using one-way ANOVA with Tukey correction. *P* values are indicated as ≤0.05*, ≤0.01**, ≤0.001***, and ≤0.0001****.

The ^13^C enrichment downstream of glucose in the lungs of wild-type animals was progressively diluted (Figure 4C). Curiously, the lungs of wild-type animals presented a lower fraction of ^13^C labeling in glycolytic intermediates and TCA cycle metabolites compared to PolG animals, suggesting that PolG animals have accelerated glucose metabolism in the normal lung and/or have increased uptake of these isotopologues from the bloodstream.

The metabolic profile in tumors markedly contrasted with that of normal lungs. The ^13^C labeling fraction of glycolytic intermediates showed notable similarity between PGKP animals on both diets and KP animals with the AA-ctr diet (Figure 4D). However, the levels of labeled downstream intermediates, particularly pyruvate, lactate, and alanine, were diminished in tumors of KP animals subjected to ser/gly starvation (Figure 4E). These findings indicate that ser/gly starvation induces adaptation to glucose metabolism in KP, but PGKP tumors have lost this metabolic flexibility.

Alterations in glucose metabolism can directly affect the synthesis of serine, which requires glycolytic intermediates and NAD+ to be synthesized. Serine M+3 is the product of the first turn of the SSP from glycolysis (Figure 4A). The labeling fraction of serine M+3 in all tissues of cancer-free PolG animals subjected to ser/gly starvation tends to be higher than that from the AA-crt diet (Figure 4F); whereas in wild-type mice, except for lung tissues, the level of serine M+3 is lower in ser/gly starvation compared to AA-crt diet (Figure 4F). However, this serine M+3 pattern was altered in NSCLC tissues. The levels of serine M+3 in KP tumors were higher than in any other tissue and condition. In comparison to KP tumors under both dietary conditions, PGKP tumors exhibited a significantly reduced ^13^C labeling fraction from glucose to serine M+3, indicating reduced utilization of glucose for serine biosynthesis. Interestingly, the bloodstream serine levels remained unaffected by serine/glycine starvation in animals with NSCLC (Figure 4G). Therefore, the decreased serine levels in PGKP tumors (Figures 3F, 4G) could be attributed to the reduced *de novo* serine synthesis from glucose. These findings support the idea that the accumulation of pathogenic mtDNA mutations and defects in the ETC creates a greater dependency on the use of glucose as a fuel source for glycolysis and suppression of serine synthesis. This metabolic defect elevates the requirement for exogenous serine and glycine to support downstream anabolic metabolism and tumor growth.

### Low NAD+/NADH levels may fail to adequately sustain both SSP and glycolysis in tumors with respiration defects

To determine if glucose use to produce serine was different in KP and PGKP TDCLs, we performed untargeted metabolomics (Figure S3A). As expected, the metabolomic data from TDCLs were similar to those from tumor tissues. The PGKP TDCLs presented a different profile compared with KP TDCLs, with the most significantly altered metabolites being intermediates of glycolysis and the SSP. Moreover, the pathway impact analysis showed that the differences between PGKP and KP TDCLs were indeed concentrated in glycolysis and serine, glycine, purine, and pyrimidine metabolism pathways (Figure S3B), suggesting that mitochondrial dysfunction in TDCLs altered glycolysis flow to the SSP.

These results, aligned with the results observed *in vivo*, suggested that PGKP tumor cells used greater levels of external glucose to carry out glycolysis and downstream metabolic pathways. Because the source of nutrients could be better evaluated and manipulated *in vitro*, we conducted a kinetic experiment extracting metabolites from TDCLs at three-time points without culture medium replacement (Figure 5A). Most glycolytic intermediates in the preparatory phase of glycolysis (Glucose-6P → Glyceraldehyde-3P) were present at higher levels in PGKP than in KP TDCLs after 24h, but the levels of these intermediates decreased over time. In contrast, the levels of these intermediates in KP TDCLs increased over time (Figure 5B). Intriguingly, the pattern of intermediates in the pay-off phase of glycolysis (3-Phosphoglycerate→Pyruvate) was significantly different. Specifically, starting from the step GAP→1,3BPG, which is also the hub to SSP. At this point, most of these intermediates were in lower levels in PGKP than in KP TDCLs. The oxidation of GAP is catalyzed by the enzyme GAPDH, which requires NAD+.^62^ Notably, GAPDH is subject to allosteric inhibition by NADH.^43,44^ In PGKP tumor cells, NADH levels are consistently elevated in PGKP compared to KP TDCLs, leading to a low NAD+/NADH ratio (Figure 5C). This disparity likely accounts for the observed glycolytic pattern.

**Figure 5.**
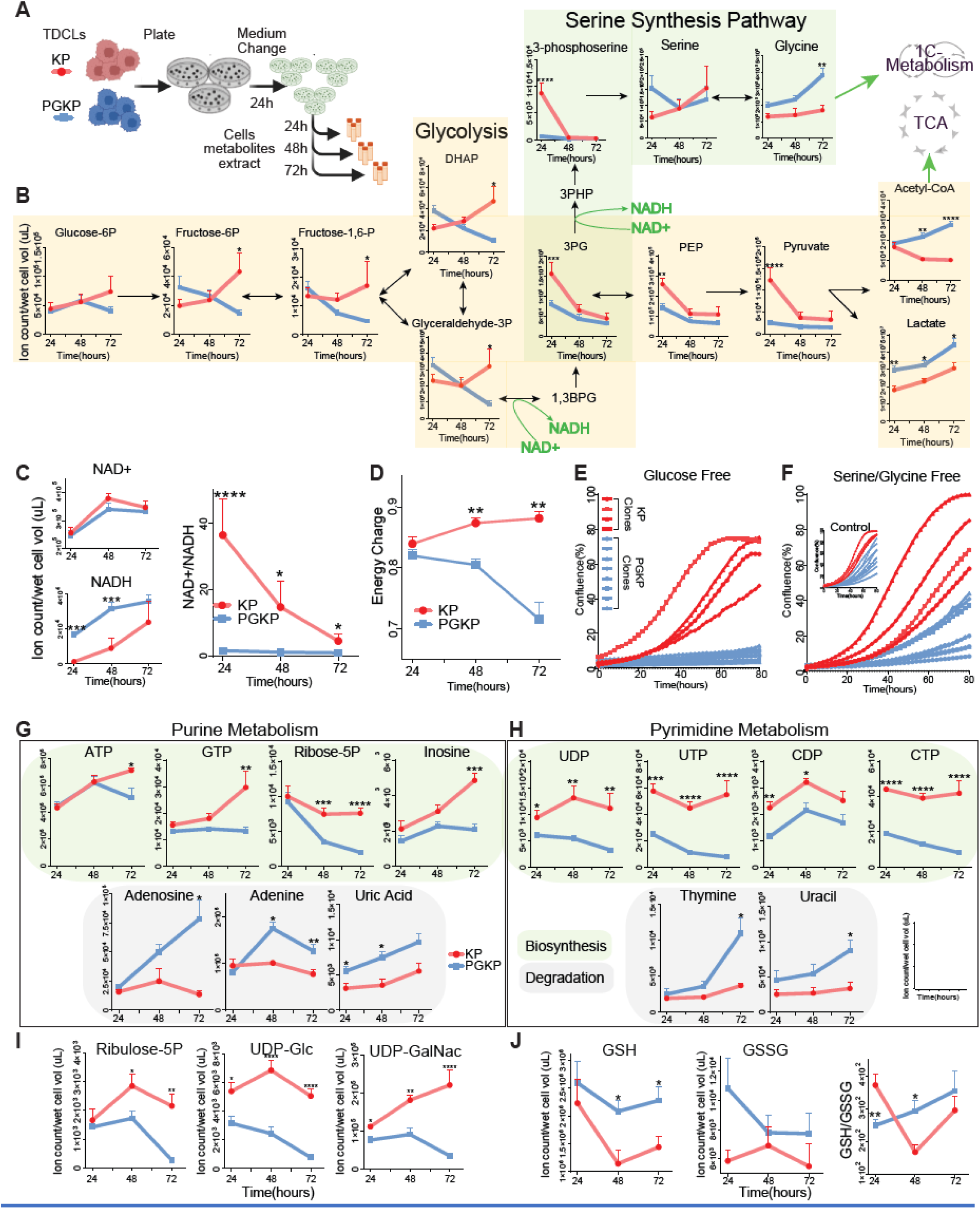
Mitochondrial respiration defects alter glycolysis and SSP, causing dependency on external glucose, serine, and glycine in NSCLC cells: **(A)** The experimental design of pool-size metabolomic kinetics. **(B)** Kinetic of key metabolic intermediates of glycolysis and SSP. Dihydroxyacetone phosphate (DHAP), 1,3-Biphosphosglycerate (1,3BPG), 3-Phosphohydroxypyruvate (3PHP), and phosphoenolpyruvate (PEP). **(C)** Kinetics of NAD+, NADH, and the resulting NAD+/NADH ratio over time. **(D)** adenylated energy charge variation over time. **(E-F)**, KP and PGKP TDCLs growth curve (Incucyte) under glucose and serine/glycine starvation. The control graph depicted in panel E is a projection of the figure 2F. **(G-H)** Kinetics of metabolic intermediates in Purine and Pyrimidine metabolisms. Green-shaded intermediates exemplify metabolites involved in nucleotide synthesis, while those in grey represent metabolic intermediates associated with nucleotide degradation. **(I)** Metabolites of the pentose phosphate pathway and sugar nucleotides. UDP-glucose (UDP-Glc). Uridine diphosphate N-acetylgalactosamine (UDP-GalNac). **(J)** GSH, GSSG levels, and their ratio variation over time. All experiments used four KP and eight PGKP-independent TDCLs. Data is shown as ± SEM. *P* was calculated using one-way ANOVA with Tukey correction and indicated as ≤0.05*, ≤0.01**, ≤0.001***, and ≤0.0001****.

Over time, there was a gradual accumulation of acetyl-CoA in PGKP TDLCs. The diminished levels of citrate in these cells indicated a limitation in the condensation of oxaloacetate with acetyl-CoA to produce citrate, potentially hindering the flow of the TCA cycle (Figure S3C). Furthermore, the levels of nearly all TCA cycle intermediates were notably lower in PGKP compared to KP TDCLs, affirming the malfunctioning state of the TCA cycle in PGKP tumor cells, as corroborated by the reduced OCR (Figure 2M). Moreover, the low NAD+/NADH ratio may further impede the proper functionality of the TCA cycle^63^, potentially affecting the fitness of PGKP tumor cells. As a result, PGKP tumor cells have defective energetics over time without replenishment of external nutrients, which ultimately was reflected in decreased energy charge (Figure. 5D),

The energetic and glycolysis alterations suggested that PGKP TDCLs may have elevated dependence on glucose as a fuel source. To test this hypothesis, we removed the glucose from the culture media of TDCLs (Figures 5E and S3D). Although the KP TDCLs had slightly reduced proliferation in the absence of glucose most of the clones reached at least 70% confluence by 80 hours. In contrast, the PGKP TDCLs showed a complete impairment in cell growth, confirming, as expected, that NSCLC cells carrying dysfunctional mitochondria are exquisitely dependent on glucose as a fuel source.

Remarkably, as observed in vivo (Figure 3D), PGKP TDCLs exhibited significantly low levels of 3-phosphoserine (Figure S3A). This finding underscores the heightened dependence of PGKP TDCLs on external sources of glucose, serine, and glycine, potentially exerting adverse effects on downstream metabolic pathways such as the SSP. Notably, the initial committed step in the SSP, which necessitates NAD+, is catalyzed by the enzyme phosphoglycerate dehydrogenase (PHGDH)^64,65^, thereby implicating a potential link between low NAD+/NADH ratio and reduced serine synthesis. To test this hypothesis, we starved TDCLs of serine and glycine (Figure 5F and S3D). Proliferation in KP TDCLs was slightly repressed by approximately 15% in serine/glycine-free medium, but the proliferation of PGKP TDCLs was decreased by almost 60%. Thus, respiration defects in NSCLC increased dependency on extracellular glucose, serine, and glycine *in vivo* and *in vitro* further exacerbated by the low NAD+/NADH that suppresses the SSP.

### Defects in respiration increase dependency on external sources of glucose, serine, and glycine to maintain GSH and nucleotide pools

The serine produced from glucose through glycolysis is one of the main sources of newly synthesized glycine, which is extensively used in anabolic metabolisms, such as 1C-metabolism^66^. Alterations in one-carbon pools can impact in nucleotide availability due to the lack of important intermediates of the methionine cycle, folate cycle, and purine and pyrimidine synthesis^18,67^.

The levels of metabolic intermediates in purine and pyrimidine synthesis in PGKP TDCLs were very different from that of KPs (Figure 5G-H). The levels of intermediates contributing to the biosynthesis of purines and pyrimidines were very low or decreased over time in PGKP TDCLs, whereas the intermediates representing the degradation in both pathways increased. Similar results could be found in other important anabolic metabolites, such as pentose phosphate pathway intermediates and nucleotide sugars (Figure 5I). In contrast, the levels of newly synthesized metabolites increased while metabolites representing degradation remained similar in KP TDCLs. These results suggested that NSCLC cells increased *de novo* synthesis of anabolic metabolites when the external nutrients were being depleted, which is normally expected. In contrast, mitochondrial respiration defects in NSCLC cells decreased the *de novo* synthesis and failed to maintain essential anabolic metabolite levels, such as the pool of nucleotides.

Another important downstream product of the SSP and synthesis of glycine is GSH. GSH helps maintain redox homeostasis and is also synthesized in response to ROS generation, which is usually increased in tumor cells^68–70^. Different from nucleotides, GSH was present at higher levels in PGKP TDCLs (Figure 5J), most likely due to ROS accumulation (Figure S3E). Mitochondrial respiration defects and suppression of the SSP would be detrimental to the tumor cell fitness due to insufficient substrate production (e.g., glycine) to synthesize GSH. GSH synthesis also requires ATP, which is also negatively impacted by tumor cell respiration defects. To test this hypothesis, we conducted an *in vitro* isotope tracing by feeding TDCLs [U-^13^C]Glucose in a complete or serine/glycine-free medium and then evaluated the glucose carbon fate in these tumor cells (Figure 6A and S4A). a As observed *in vivo*, KP TDCLs used more glucose than PGKPs in regular media to synthesize serine, but *in vitro*, we could clearly see that glycine was also affected (Figure 6B and S4B-C). Moreover, the synthesis of GSH from glucose also impacted PGKP tumor cells (Figure 6B and Figure S4D), showing that mitochondrial dysfunction impaired the use of glucose to produce glycine and GSH. The production of these metabolites using glucose carbons was increased in serine/glycine-starved KP tumor cells, which partially rescued the internal levels of these amino acids and the levels of GSH. These results were not observed in PGKP tumor cells since the total serine/glycine levels were very low, and the M+2 (from glycine or glutamate alone) and M+4 (from glycine+glutamate) isotopologues of GSH barely changed; as a result, the total levels of GSH could not be rescued (Figure 6B and S4D).

**Figure 6.**
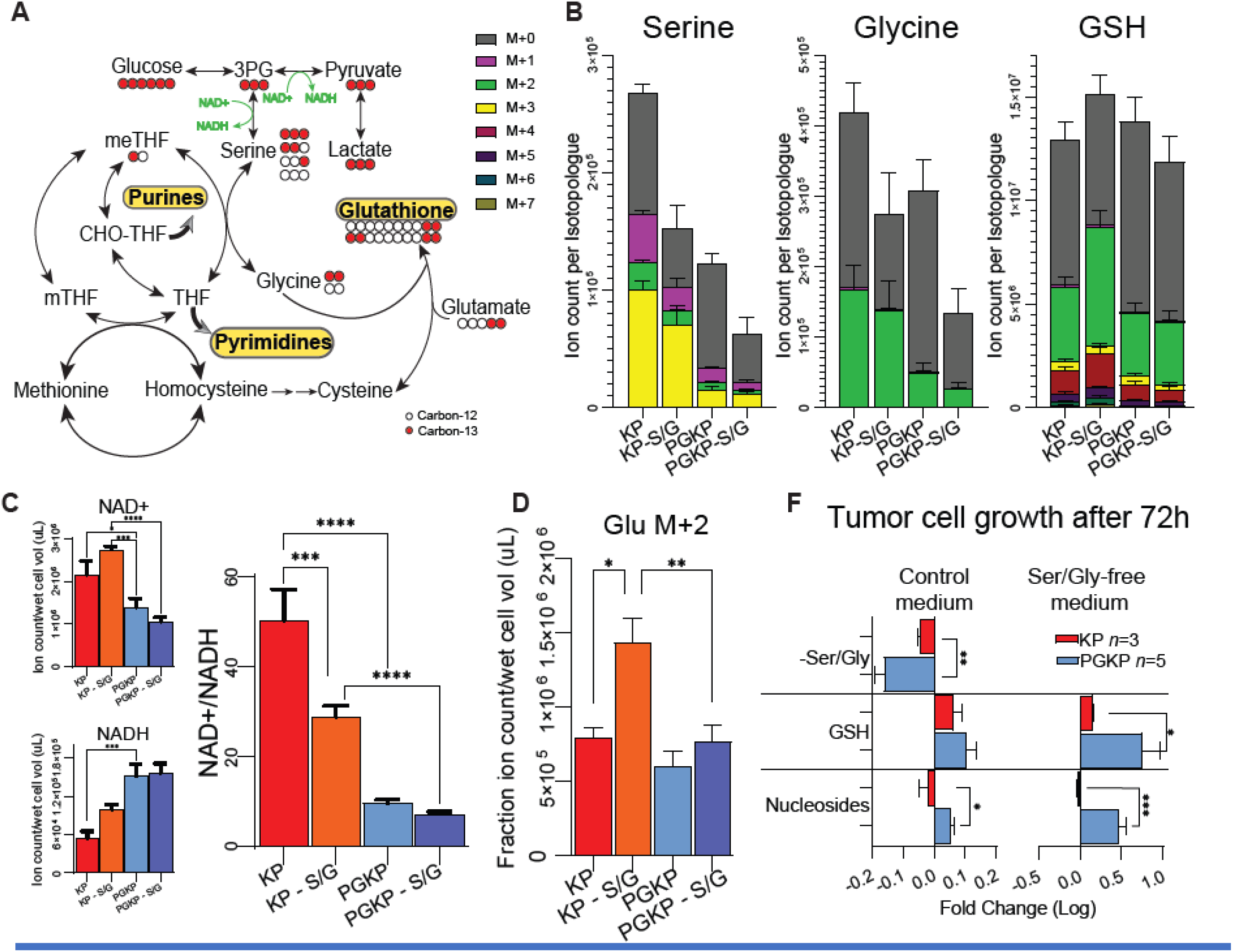
NSCLC cells rewire glucose metabolism during serine/glycine-starvation to feed SSP and 1C-metabolism: **(A)** Schematic of carbons flux from glucose to serine, glycine, and GSH using the glycolysis→SSP→1C-metabolism axes. M+0 to M+7 represents the number of ^13^C present in the isotopologue. **(B)** Pool size levels of each isotopologue of serine, glycine, and GSH (labeled fractions + M+0) in KP (n=3 clones) and PGKP (n=5 clones) TDCLs cultivated in both control and serine/glycine-starved medium. The schematic experimental design and statistical analysis details are outlined in Figure S4. **(C)** NAD+ and NADH levels and resulting NAD+/NADH ratio of KP and PGKP TDCLs in complete and ser/gly-starved medium. **(D)** Glutamate M+2 pool size. **(H)** Growth of KP and PGKP TDCLs cultivated for 72 hours in both control and serine/glycine-starved medium, with supplementation (or without) of GSH (5mM) and nucleosides. The graph depicts the fold change of the control group compared to the supplemented group. Data is shown as ± SEM. For all figures, the *P* values were determined using one-way ANOVA with Tukey correction. *P* values are indicated as ≤0.05*, ≤0.01**, ≤0.001***, and ≤0.0001****.

Once more, the NAD+/NADH ratio emerges as the critical factor in this process (Figure 6C). NAD+ levels were lower in PGKP than in KP tumor cells and further declined when these TDCLs were deprived of serine/glycine. This supports the hypothesis that NSCLC cells with respiratory defects have an NAD+/NADH imbalance exacerbating redox activity impairment, particularly during serine/glycine starvation.

Additionally, under serine/glycine starvation, KP TDCLs exhibited an accelerated TCA cycle without a corresponding increase in carbon consumption from glucose. This was evident in the elevated total levels of most TCA cycle intermediates (ions count), while the labeled fractions of these metabolites remained unaltered (constant M+0) (Figure 6D and Figure S5). Moreover, as glutamate is the most abundant TCA cycle-associated metabolite, it is possible to infer TCA flux by checking M+2 glutamate levels^7^. This result suggests TCA cycle acceleration in serine/glycine-starved KP tumor cells (Figure 6D and S5). In sum, this process ensures that NSCLC cells under nutrient-deprived conditions maintain energetic and anabolic homeostasis by maintaining appropriate NAD+/NADH levels, which cannot be achieved by tumor cells with defective mitochondrial respiration, as observed in PGKP TDCLs.

To investigate whether we could rescue the growth of PGKP TDCLs subjected to serine/glycine starvation, we provided the depleted downstream metabolites through supplementation with GSH and nucleosides. GSH and nucleoside supplementation significantly rescued the serine/glycine-starved PGKP TDCL growth compared with KP, showing that these metabolites are important fates for external serine/glycine (Figure 6F). Curiously, GSH supplementation also improved KP TDCL growth under serine/glycine starvation, but to a lesser extent. Additionally, the GSH levels in serine/glycine-starved KP tumor cells exhibit a tendency to rise, surpassing even those in KP cells cultured in a complete medium (Figure 6B). Moreover, their reliance on glucose for GSH synthesis notably escalates compared to any other group (Figure 6B higher enriched GSH and S4D lower M+0 fraction). This data suggests that NSCLC cells also benefited from external serine and glycine to keep the optimum levels of GSH, but mitochondrial dysfunction further increases this need. To establish the connection between the availability of external serine/glycine, accumulation of ROS, and GSH levels, we measured ROS levels under control and serine/glycine-starved conditions (Figure S6). Indeed, under serine/glycine starvation, KP tumor cells increased ROS levels after 72h. However, the same was not observed in PGKP tumor cells, presumably due to their already elevated basal levels of ROS. These data suggest that KP NSCLC controls ROS levels using external serine/glycine to synthesize GSH, switching to SSP under starvation of these amino acids as a compensatory mechanism. Moreover, serine/glycine starvation, most likely due to an accelerated TCA cycle (as showed elsewhere^17^ and here in KP tumor cells) or defects in mitochondrial respiration generates even more ROS to compensate for the lack of glucose that might be deviated to SSP and, thereby, nucleotides and GSH synthesis.^17^

## DISCUSSION

Mitochondria play a major role in many cellular functions such as energy generation, biosynthesis (e.g., amino acids, iron-sulfur clusters, heme), redox control, differentiation, and signaling.^22,71,72^ The role of mitochondria in cancer is not well understood, nor is the role of specific aspects of mitochondrial function.^73,74^ A subset of rare tumors mostly arising in the kidney are characterized by pathogenic mtDNA mutations.^75–77^ In these tumors, the pathogenic mtDNA mutations are often homoplastic and appear to be founder events, suggesting that in rare cases, tumorigenesis can be initiated by loss of mitochondrial function. Other good examples are thyroid and colon cancers since these tumors carry a high mtDNA mutation burden and accumulate dysfunctional mitochondria.^4,28,78^ However, in all these cases, the mechanistic role of mtDNA mutations is poorly understood.

Most cancers, however, appear to preserve mitochondrial function by selecting against the accumulation of pathogenetic mtDNA mutations.^28,79^ There are also examples where tumor cells engineered to have mtDNA eliminated reacquire mtDNA from their host.^80^ Indeed, knocking out the essential mitochondrial transcription factor *Tfam* results in impaired tumor growth.^74^ Given the role of mitochondria in energy generation, numerous biosynthetic processes, redox control, and signaling, it might be expected that most tumors derive some benefit from maintaining mitochondrial function. This further suggests that there are distinct roles for mitochondria in different cancer types, with most needing them and other rare settings where loss of mitochondrial function may initiate tumorigenesis. We sought to address the question of the tumors that benefit from mitochondrial function; what aspects of their function are important in cancer?

Tumor growth has been found to require the ETC to oxidize ubiquinol, which is essential to drive the oxidative TCA cycle, and dihydroorotate dehydrogenase (DHODH) activity is essential for de novo pyrimidine synthesis.^81,82^ Additionally, restoration of NAD+ levels in respiration defective tumor cells through the expression of *Lactobacillus brevis* NADH oxidase (*LbNOX*), which mimics complex III activity, fails to rescue tumor cell growth^40^. These findings, coupled with our results, imply that maintaining a balanced NAD+/NADH ratio, rather than simply elevating NAD+ levels, is crucial for tumor cell metabolic fitness.

We report here that engineering NSCLCs to accumulate pathogenetic mtDNA mutations through the introduction of a *PolG*^*D256A*/*D256A*^ mutator allele significantly extends mouse survival, indicating that they rely on mitochondrial function. These *PolG*^*D256A*/*D256A*^ mutator tumors and TDCLs have a cell-intrinsic defect manifested by decreased proliferation and viability contributing to defective tumor growth. To determine the cause of these proliferative and survival defects, we examined the metabolic alterations in the *PolG*^*D256A*/*D256A*^ mutator tumors and TDCLs. While the accumulation of mtDNA mutations in NSCLC caused the expected glucose dependency and glycolytic metabolism with impaired respiration, it also resulted in a decrease in metabolites downstream in the SSP that feed the synthesis of glutathione and nucleotides. Indeed, *PolG*^*D256A*/*D256A*^ mutator tumor cells were more dependent on exogenous serine and glycine for survival, and supplementation with nucleosides and glutathione downstream in the SSP partially rescued growth and survival. Finally, while a serine and glycine-free diet had no effect on mice with NSCLC, it further impaired tumorigenesis and extended survival in mice bearing NSCLC with the *PolG*^*D256A*/*D256A*^ mutator allele and pathogenic mtDNA mutations. This indicates that impairing mitochondrial function creates a metabolic liability, leading to an increased reliance on glucose due to respiratory impairment. This respiratory impairment causes the accumulation of NADH and an imbalanced NAD+/NADH ratio, which is crucial for maintaining the redox balance necessary for both glycolysis and the SSP, affecting the availability of 1C units crucial for key biosynthetic reactions such as purine synthesis.^83,84^ Since serine serves as the primary source of 1C units, NSCLC cells with respiratory deficiencies develop a dependence on external serine and glycine sources to sustain robust tumor cell growth. Moreover, the impairment in the SSP limits the ability to produce nucleosides and GSH, which is likely responsible for redox stress, reduced growth, and survival. Compensating for limiting glucose to fuel both glycolysis and the SSP with a high carbohydrate diet reversed the survival advantage of the *PolG*^*D256A*/*D256A*^ mutator NSCLC. Thus, NSCLC preserves mtDNA to prevent the metabolic liability associated with respiration defects and exclusive dependency on glucose as a fuel source that limits serine utilization for biosynthesis.

How respiration defects lead to suppression of the SSP is likely related to elevation of NADH and lowering of the NAD+/NADH ratio. Elevated NADH levels can impair glycolysis, as Glyceraldehyde-3-Phosphate Dehydrogenase (GAPDH) is allosterically inhibited by NADH.^43,44^ This inhibition not only affects the rate of glycolysis but also impacts the availability of crucial glycolytic substrates such as 3-phosphoglycerate (3PG).^45^ Given that a significant proportion of glycolytic 3PG is rerouted into the serine and glycine biosynthetic pathway,^46^ an imbalanced NAD+/NADH ratio can directly influence the flux of the serine synthesis pathway.

These findings raise a few important points. First, as metabolism is distinct across different cancers, do other cancers similarly depend on the preservation of mtDNA and mitochondrial function to fuel the SSP? Does the upregulation of the SSP alone sustain NSCLC growth and survival, or do these tumor cells essentially depend on a balanced NAD+/NADH ratio to sustain optimal growth? Upregulation of the SSP through amplification or upregulation of *PHGDH* expression is seen in some cancers, which illustrates the functional importance of this pathway,^85–87^ but whether this is in addition to the preservation of mitochondrial function or an adaptation to mitochondrial deficiency is unknown. Second, while targeting metabolism is a mainstay of chemotherapy, our findings indicate that a deeper understanding of cancer metabolism can reveal novel metabolic dependencies that can be targeted. Third, we show that a deep understanding of cancer metabolism can also reveal metabolic vulnerabilities that can be targeted by dietary manipulation. While the concept of diet as therapy is not new, determining what diet to use in specific cancer settings for both prevention and treatment is worth further exploration. New therapeutic approaches could include combining the mitochondrial Complex I inhibitor Metformin with a serine and glycine-free diet in NSCLC or treating rare tumors with pathogenic mtDNA mutations (as evidenced by the ND6KP model) with dietary serine and glycine deprivation or SSP inhibitors.

## Acknowledgments

This work was supported by R01 CA243547 to EW, ECL, and SG and RO1 CA259635 to DCW. Duncan and Nancy MacMillan Center of Excellence in Cancer Immunology and Metabolism to CSH, EW, SG and CSC. The Biospecimen Repository and Histopathology Service Shared Resource from the Cancer Institute of New Jersey provided all the specimens and associated services (P30CA072720-5919); Biometrics Shared Resource (P30CA072720-5918); Metabolomics Shared Resource (P30CA072720-5923). Biomedical Informatics Shared Resources (P30CA072720-5917). We thank Jennifer Hostettler from the Medical Writing Services of Rutgers Cancer Institute of New Jersey and Li-Kuo Su from the Scientific Editors for assistance in editing this manuscript.

## Author Contribution

ECL performed or participated in all experimental work, PGKP and KPND6 GEMM generation and characterization, data analysis, data interpretation, and wrote the paper. EW is the leading investigator who conceived the project, supervised research and edited the paper. JR, JYG, SG, and EL provided intellectual input for the experiments, data analysis, and data interpretation. AS, MI, and MGJ assisted with mouse experiments, data analysis, and interpretation. ZH performed jugular vein catheterization and assisted with KPND6 generation. ZH and PM assisted with mouse colony management and mouse experiments. VB assisted with OCR and ECAR experiments. FS and CC processed, interpreted, and analyzed mtDNA and WES sequencing data. WW assisted with in vivo isotope tracing experiments. XS performed metabolomic processing and assisted with data analysis. DCW contributed advice on mouse models of mitochondrial disease.CSH and all other authors participated in the paper editing process.

**Figure S1.**
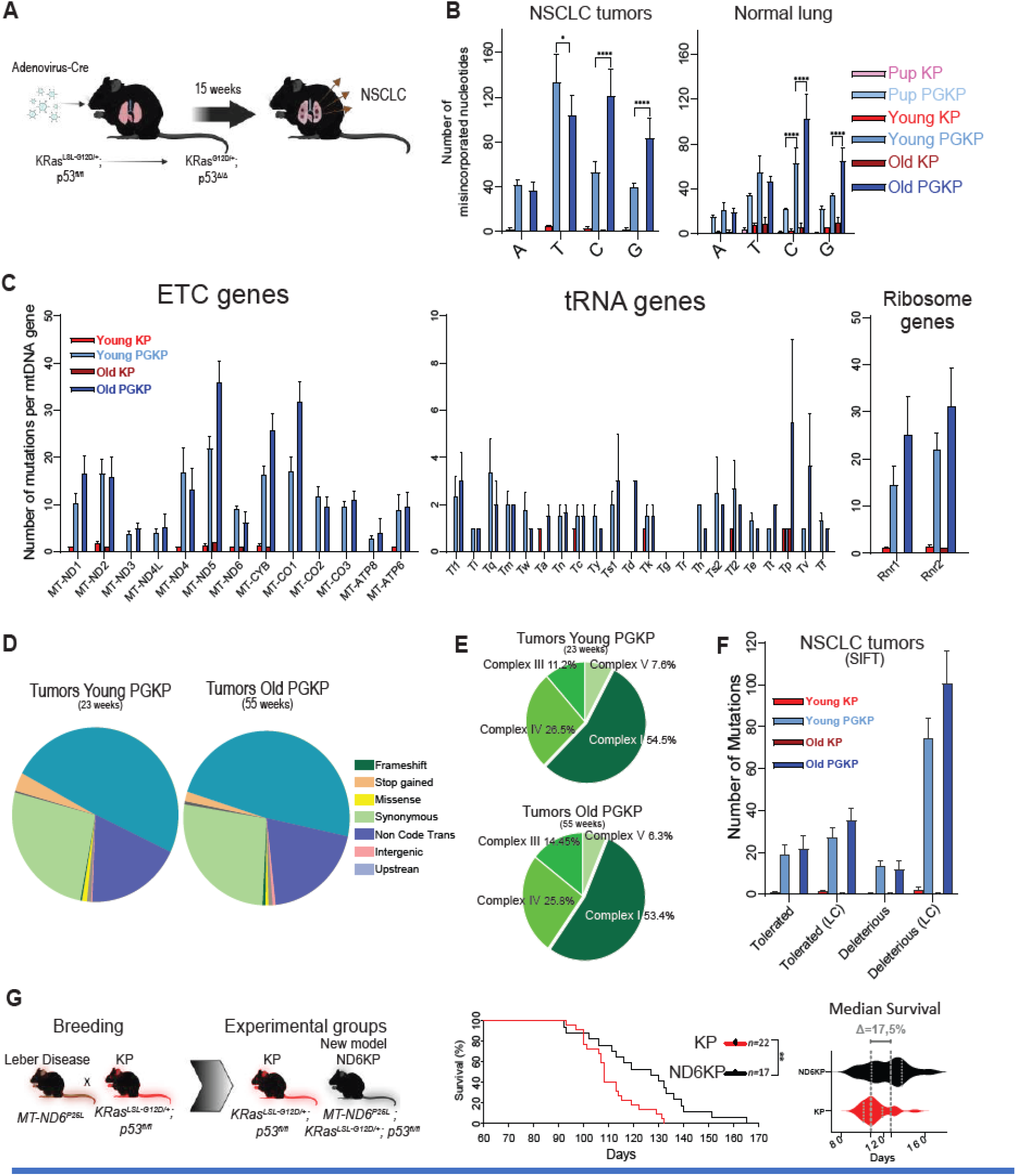
PGKP mice accumulate pathogenic mtDNA mutation burden over time: **(A)** Diagram depicting NSCLC initiation via adenovirus-Cre inhalation. Tissue collection occurs 15 weeks post-tumor initiation unless otherwise specified, in accordance with the experimental design. **(B)** mtDNA misincorporated nucleotides in NSCLC tumors and normal lung tissues. Normal lungs are extracted from PolG and WT animals. **(C)** Number of mutations per mtDNA gene. **(D)** The profile of the types of mtDNA alterations found in young and old PGKP tumors. **(E)** Percentage of mtDNA mutations found per each ETC complex in young and old PGKP NSCLC tumors. **(F)** Sort Intolerant From Tolerant (SIFT) number of mtDNA mutations predicted as pathogenic (LC= low confidence). **(G)** The diagram illustrates the generation of the ND6KP mouse model, their survival curve, and median survival after NSCLC initiation. Data is shown as ± SEM. In panels B and F, only the statistical analysis comparing young and old PGKPs is presented. The comparisons between PGKP and KP animals yielded *p*-values all ≤0.0001. Log-rank (Mantel-Cox) calculated P values for the survival curve. The *P* values were determined using one-way ANOVA with Tukey correction. The *P* values are indicated as ≤0.05*, ≤0.01**, ≤0.001***, and ≤0.0001****.

**Figure S2.**
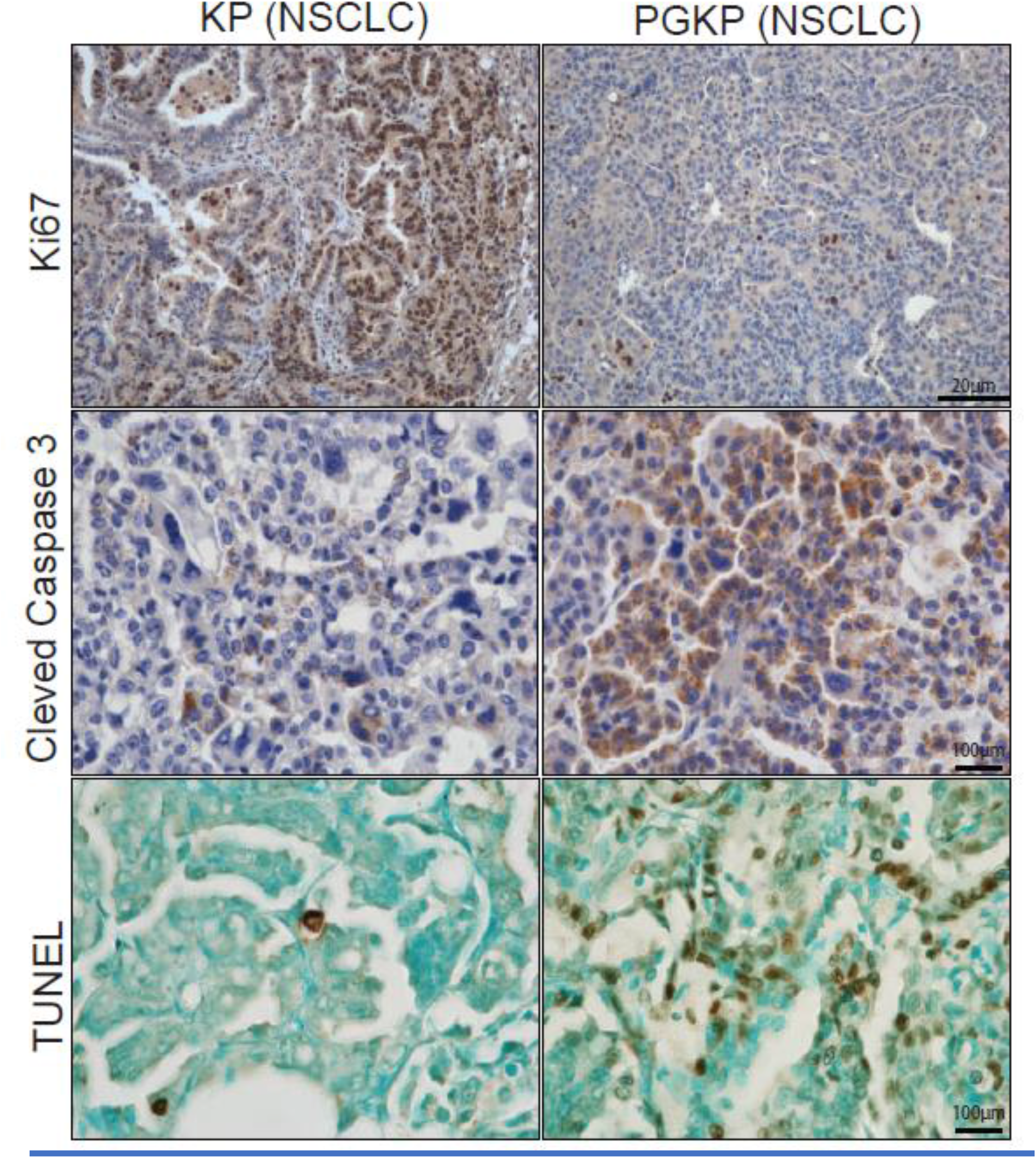
PGKP tumors exhibit reduced viability and proliferation compared to KPs. Representative pictures of IHC for Ki67 and cleaved caspase 3 and TUNEL assay of young PGKP and KP performed in tumors harvested 15 weeks post NSCLC initiation.

**Figure S3.**
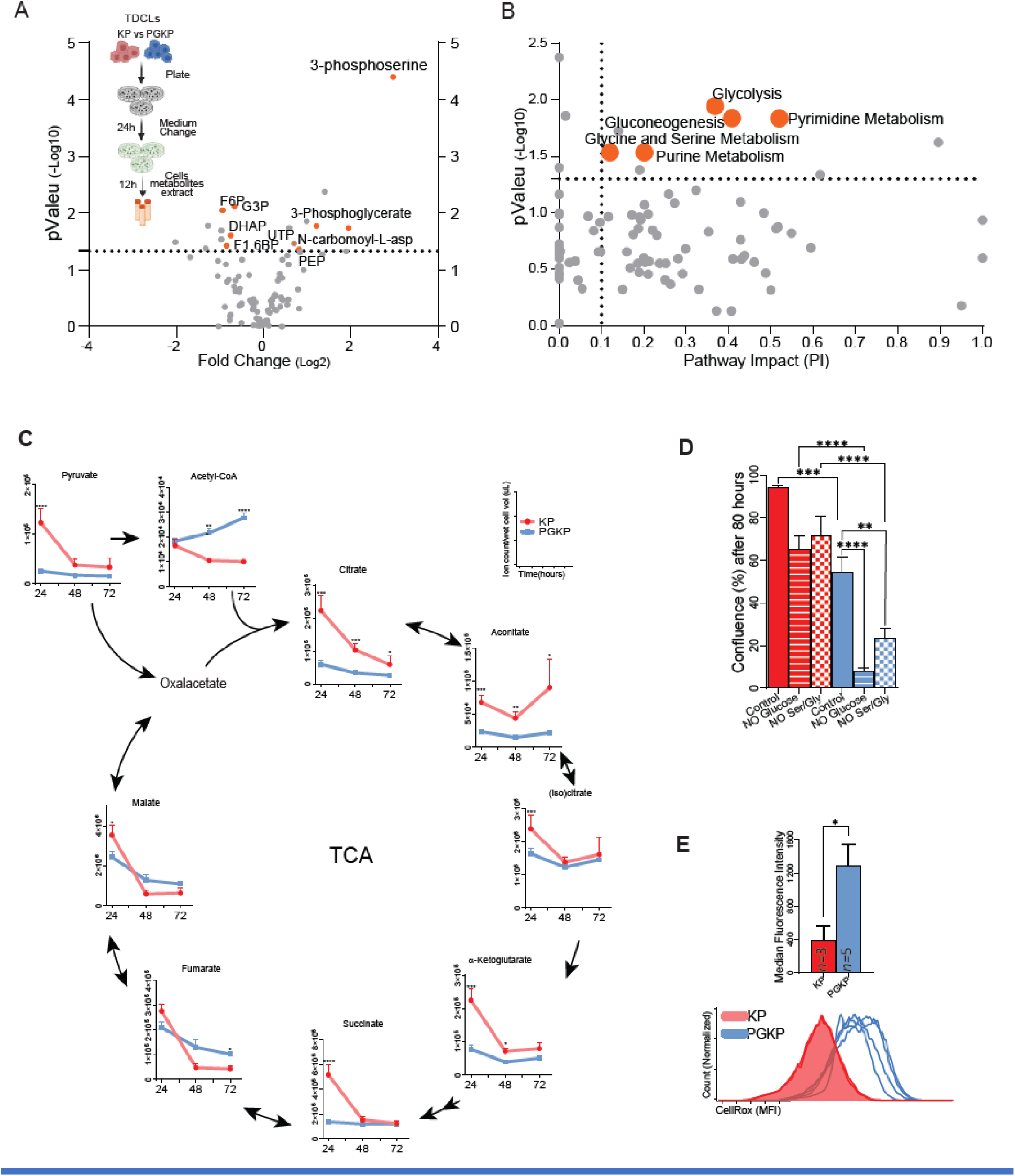
PGKP tumor cells present lower levels of TCA cycle intermediates and high ROS levels: **(A)** Schematic of pool-size metabolomics with the corresponding volcano plot illustrating the metabolic profile of KP and PGKP TDCLs. **(B)** The pathway impact analysis contrasts KP and PGKP TDCLs metabolic alterations in energetic (glycolysis) and anabolic metabolism (Pyrimidine/Purine and serine/glycine metabolism). **(C)** Kinetic of intermediates of TCA cycle contrasting KP and PGKP TDCLs metabolism. The experimental design is presented in Figure 5A. **(D)** Confluence of TDCLs at the end of 80h. **(E)** Flow cytometry presents the basal ROS levels (median fluorescence) in KP and PGKP TDCLs. Data is shown as ± SEM. The *P* values of C and D were determined using one-way ANOVA with Tukey correction. The *P* values of E were determined using two-tailed t-tests and indicated as ≤0.05*, ≤0.01**, ≤0.001***, and ≤0.0001****.

**Figure S4.**
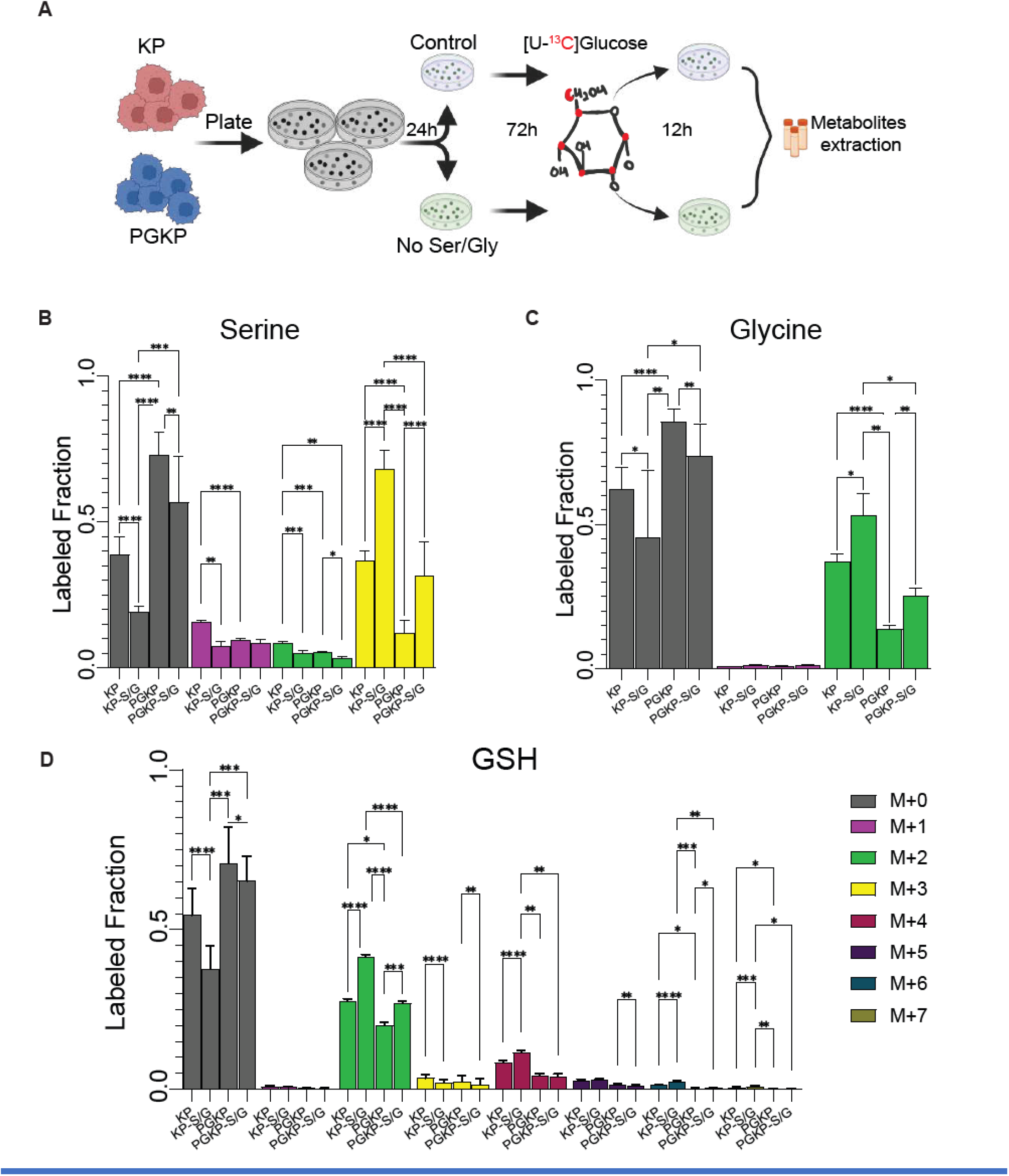
Mitochondrial respiration dysfunction, associated with serine/glycine starvation, leads to serine, glycine, and GSH synthesis deficiencies in NSCLC cells. **(A)** Schematic experimental design of pool-size metabolomics under 12 hours of [U-^13^C]-glucose labeling. KP and PGKP TDCLs were previously cultured in a control or serine/glycine-free medium for 72 hours. Subsequently, the medium was changed to the equivalent condition, and [U-^13^C]-glucose was administered for 12 hours of incubation following metabolite extraction. **(B-D)** Statistical analysis per isotopologue fraction (%) of serine, glycine, and GSH. Data is shown as ± SEM. The *P* values were determined using one-way ANOVA with Tukey correction and indicated as ≤0.05*, ≤0.01**, ≤0.001***, and ≤0.0001****.

**Figure S5.**
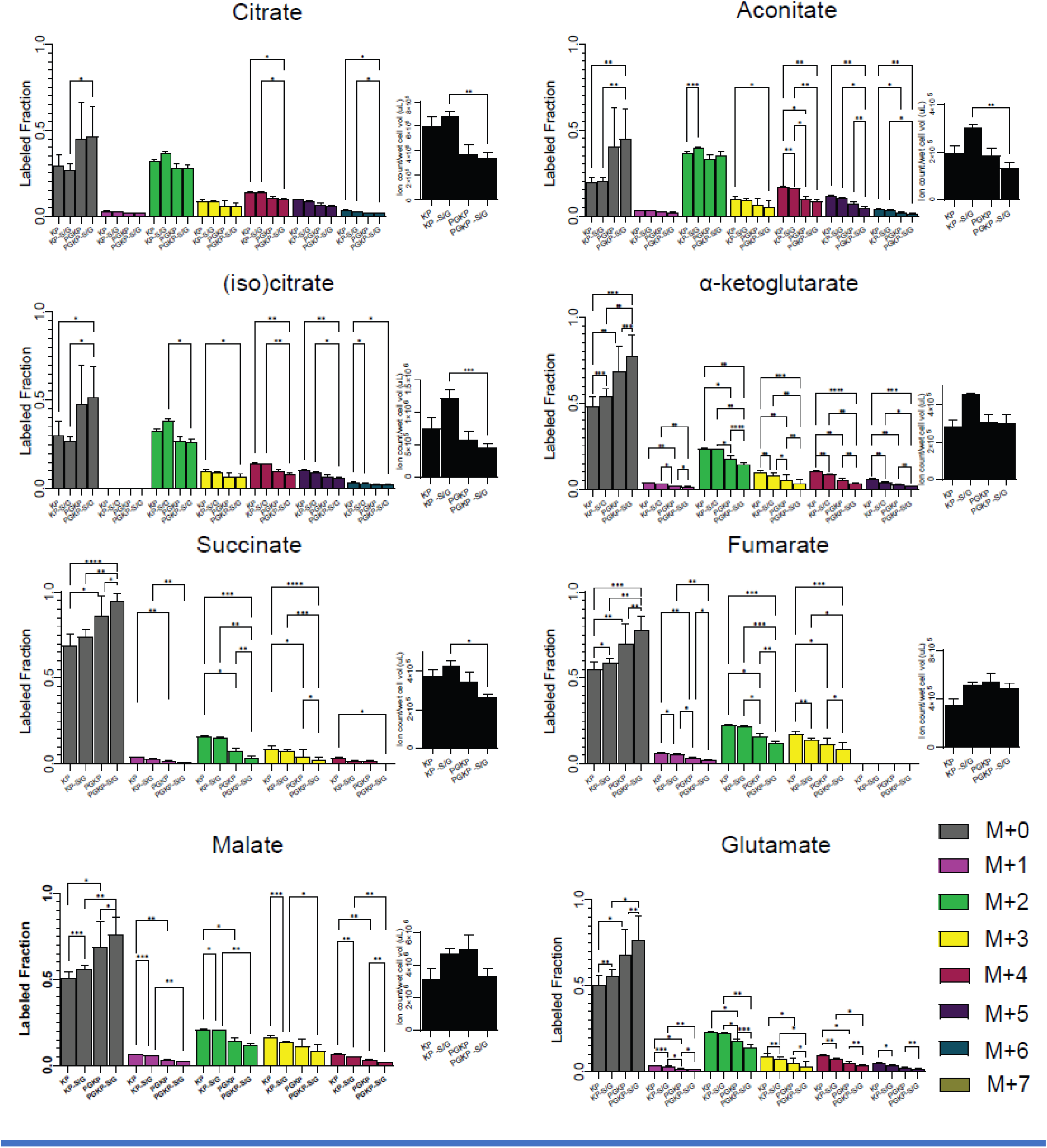
Serine/glycine starvation causes TCA cycle acceleration in NSCLC tumor cells to compensate for the lack of carbons provided by glucose: Statistical analysis per isotopologue fraction (%) and total pool size of TCA cycle intermediates. Data is presented as ± SEM. The *P* values were determined using one-way ANOVA with Tukey correction and indicated as ≤0.05*, ≤0.01**, ≤0.001***, and ≤0.0001****.

**Figure S6:**
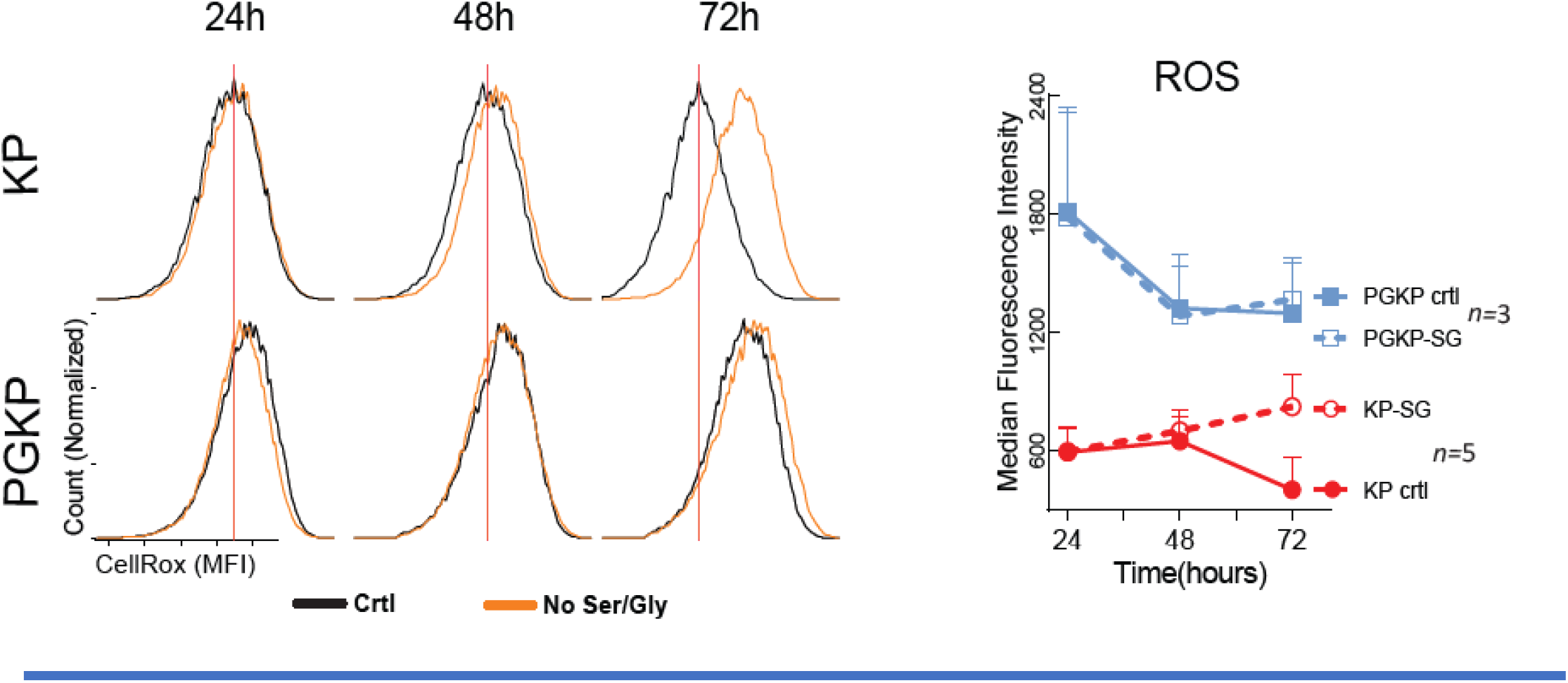
Serine/glycine starvation causes increased ROS levels in NSCLC cells: Flow cytometry measuring ROS accumulation over time in KP and PGKP TDCLs under control or serine/glycine-free medium. All experiments used three KP and five PGKP-independent TDCLs. The figure is an illustrative representation of one clone per genotype. The red lines indicate the median fluorescence intensity (MFI= ROS levels) of KP TDCLs in the control medium.

## METHODS

### Genetically engineered mice models

All animal care procedures, experiments, and maintenance complied with the Rutgers University Institutional Animal Care and Use Committee (IACUC). The KP mouse model (The Jackson Laboratory)^32^ was cross-bred with the PolG mouse model (The Jackson Laboratory reference) to generate the PGKP mouse model. The ND6 mouse model (generously provided by Dr. Douglas C. Wallace)^52^ was cross-bred with the KP mouse model to generate the ND6KP mouse model. NSCLC is initiated through intranasal infection with adenovirus-Cre 4×10^7^ pfu per mouse at 8-10 weeks (young) and ten months old (old).

For survival analysis, the animals were infected and monitored weekly until the 10th week. After this period, they were monitored daily until they reached the humane endpoint (body condition score 2). The animals’ weight and food intake were measured weekly (when indicated) until the end of the experiment.

### Purified diets

The animals were weighed one week before tumor induction. In the same period, special food was provided and kept until the end of the experiment. The food intake and animal weight were taken weekly. The special diets were obtained from Envigo according to the product code: Serine/Glycine-free diet (TD.180296), Amino acid defined (AA-crt) diet (TD.01084), and 70% High Carbohydrate diet (TD.220722).

### Blood glucose measurement

Fresh blood was collected from the animals’ tails and immediately submitted to measurement using FreeStyle Optimum Neo glucometer (Abbott Laboratories) to determine blood glucose levels. The blood was always collected between 1-3 p.m. in *ad libitum* conditions.

### Tumor burden quantification

For tumor area measurements, H&E-stained lung samples were imaged at Rutgers Cancer Institute of New Jersey Biomedical Informatics shared resource using an Olympus VS120 whole-slide scanner at 20X magnification. The image analysis protocol was custom-developed on the Visiopharm image analysis platform to identify tissue area and number and compute tumor burden based on semiautomatically detected tumors as described previously.^55^For tumor number, the prepared H&E-stained lung sections were analyzed using a Nikon Eclipse 80i microscope.

### Cell lines generation

All KPs and PGKPs TDCLs were generated in-house directly from their corresponding mouse tumors. Animals were infected at 8-10 weeks old (young animals) or 10 months old (old animals) and kept for 15 weeks. After this period, the tumors were harvested, and each clone was generated from a different lung lobe to ensure generation of independent clones. The TDCLs were cultivated in RPMI with 10% fetal bovine serum and 1% PenStrep solution. Additionally, serine/glycine-free RPMI (Teknova) was used as indicated.

### Cell proliferation and viability assays

TDCLs were plated 5 × 10^4^ cells per well in 6-well plates in RPMI conditions for 24h. The next day, the medium was aspirated, washed with PBS, and changed for complete RPMI or serine/glycine-free RPMI plus the indicated treatments: 5mM of reduced glutathione (Sigma) or nucleoside mix (EmbryoMax®Nucleosides 100x, Millipore). The cells were kept without medium change for 72h, harvested, and then counted in triplicate using a Vi-CELL™XR cell counter and viability (Trypan blue) analyzer system with a cell viability assay using a flow cytometer (Beckman Coulter). To generate Incucyte growth curves, TDCLs were plated 5 × 10^4^ cells per well in 12-well plates in complete, glucose-free, or serine/glycine-free RPMI conditions for 80h. For viability assays using Trypan blue in Zcounter, the KP (n=4) and PGKP (n=8) TDCLs clones were tested for 13 passages twice in a regular RPIM medium. All experiments and conditions were repeated at least twice. All experiments used at least three independent TDCLs clones per genotype.

### Growth of TDCLs in mice

C57Bl/6J mice (The Jackson Laboratory) were injected (subcutaneously) with 10^5^ cells per animal flank. The tumors were measured at least thrice weekly until the animal reached the humane endpoint as determined by the protocol (<1000mm^3^ determined by *π//6 x L x W x H*).

### WES and mtDNA genome sequencing

Total DNA from tissues (tumors n=5, normal lung tissues n=3, and normal pup lung tissue n=2 from each genotype and age group) and TDCLs (KP n=4 and PGKP n=8 from each age group) were isolated. Library preparation was carried out using the Agilent SureSe-lectXT mouse mitochondrial custom enrichment protocol (Agilent Technologies). Each pooled sample was clustered and sequenced on an Illumina MiSeq instrument (Novogene Corp.) using two 150-base-pair (bp) paired-end reads and∼2.6 million reads per sample.

### mtDNA genome and whole exome sequencing data analysis

The analysis of mtDNA genome raw fastq files used FastQC v0.11.9^88^ for quality checks and Trimmomatic v0.40^89^ for read trimming. To align the reads to the reference genome GRCm38.p6, we employed bwa (Burrow-Wheeler Aligner) v0.7.17-r1188^90^. The alignment data were then evaluated using Qualimap v2.3^91^. We used samtools v1.3.1 to index and sort the resulting bam files for further processing. Duplicate marking was carried out using web-based gatk v4.1.8 (https://gatk.broadinstitute.org). To identify somatic SNVs and indels, we utilized gatk mutect2 with mitochondrial mode. To ensure the quality of the variants, we applied specific filters based on allele fractions (AF)>0.01, approximate read depth (DP)>2000, and the log 10 likelihood ratio score of variant existence (TLOD)>10. For variant annotation, we employed VEP (Variant Effect Predictor)^92^. IGV v2.8.2, R, and Python were used for data visualization.

A similar pre-processing pipeline for whole exome sequencing was employed. However, for somatic single nucleotide variants (SNVs) and indel calling, the gatk mutect2 tool in tumor-normal mode was applied. Variant annotation was performed using SnpEff v4_3t ^93^. For post-processing, we applied the following filter settings: SNVs and indels were retained if they had a variant allele frequency of ≥10%, a minimum read coverage of 10x in both the tumor and normal samples at that specific position, a minimum of three reads supporting the variant allele in the tumor sample, and no reads supporting the variant allele in the matched normal tissue. To detect copy number variants, we employed CopyrightR v1.3^94^. We also utilized R and Python to identify cancer-drive genes and visualize the data.

### Histology, TUNEL and IHC

Lungs with tumors from each experimental group were harvested after 15 weeks of NSCLC initiation (except for the tumor initiation experiment that was harvested after six weeks), fixed in 10% buffer formalin solution overnight, and then transferred to 70% ethanol for paraffin-embedded sections. Tissue sections were deparaffinized, rehydrated, and boiled for 45 min in 10 mM pH 6 citrate buffer. All Paraffin-embedded blocks were submitted to stained with hematoxylin-eosin (H&E) for quality check. Blank slides were used for IHC preparation with antibodies for Ki67 (Abcam, ab15580) and cleaved caspase 3 (Cell Signaling, 9661S). Paraffin-embedded slides were also used for the TUNEL assay and processed according to manufacturer instructions (Abcam, TUNEL Assay Kit - HRP-DAB ab206386). The prepared sections were analyzed using a Nikon Eclipse 80i microscope. ImageJ was used to extract optical density (CC3) and positive nuclei count (Ki67 and TUNEL) for quantification. Each experiment used at least four animals and ten images per animal. Each image corresponds to a different lung tumor.

### Assessment of oxygen consumption rate (OCR) and extracellular acidification ratio (ECAR measurement)

OCR in KP and PGKP TDCLs OCR was measured using a Seahorse Biosciences extracellular flux analyzer (XF24) as described previously.^95^ Briefly, cells were seeded at 10^4^ cells per well in the XF24 plates overnight before XF assay. Real-time OCR measurements were performed in RPMI, HBSS, and HBSS with 2 mM glutamine or 1 mM dimethyl-α-KG for 3 h, and measurements were taken every 15 min. Relative OCR (percentage) was normalized to the 0-min time point. ECAR and glycolysis rates were also measured using a Seahorse XF24, as previously described.^95^ Briefly, cells were seeded at 10^4^ cells per well in the XF24 plates overnight and then subjected to XF assays according to the manufacturer’s instructions to determine basal glycolysis, glycolytic levels, glycolytic capacity, and glycolytic reserve.

### Flow cytometry

Cells were plated in 6 well plates and proceeded as indicated by each experiment. All samples were run in the Attune NxT Blue-Red-Violet (Thermo Fisher), collecting 3×10^3^ events in at least three biological duplicates (independent clones) and in at least two independent experiments. MytoTracker green (Thermo) was used to measure mitochondria mass; MytoTracker red was used to measure mitochondria membrane potential according to manufacture specifications. CellROX™ Green Flow Cytometry Assay Kit (Thermo) was used to quantify the accumulation of ROS according to the manufacturer’s instructions. The raw data were treated and analyzed by FlowJo® V10.2. to extract the Median Fluorescence Intensity (MFI) per sample.

### Extraction of polar metabolites from TDCLs

TDCLs were plated in 6 multiwell plates (Corning) and incubated (37° C in 5% CO_2_) for 24h hours. After this period, the medium was replaced and cells were further incubated and extracted at the indicated time points. Three wells were used for the metabolite extraction, and the other three were harvested using trypsin 0.25% to measure the wet cell volume using a PCV-packed cell volume tube (TPP). The metabolite extraction was performed by washing the wells twice with cold PBS, and 400 µL of 40:40:20 buffer (Methanol: Acetonitrile: Water) with 0.05% formic acid was added to each well. The plate was allowed to rest on ice for 5 minutes, and then the cells and buffer were scraped. The samples were placed in a 1.5 mL microtube with 22 µL of 15% NH_4_HCO_3_ vortexed and centrifuged for 10 min at 15,000g at 4° C. 380 µL of the supernatant were collected and stored in −80° C freezer until analysis by LC-MS. For tracing, cells were incubated with 2g/L of [U-^13^C]D-glucose (Cambridge Isotope Laboratories) for 12h before metabolite extraction.

For solid tissues, 20-30 mg were weighed and added to a 2 mL round-tipped microtube with a – 80° C cold Yttria Grinding Ball per tube. The tissues were pulverized in CryoMill (Retsch) following alternating three cycles at 5 Hz for 2 min with three cycles at 25 Hz for 2 min. Buffer was added to each 2 mL microtube (sample weight x 40)/2 volume of the buffer) 40:40:20 buffer with 0.5% formic acid, samples were vigorously vortexed and incubated on ice for 10 minutes, vortexed, and incubated for an additional 10 min. After the samples were centrifuged for 10 min at 16,000g at 4° C, the supernatant A was collected and saved, and the pellets were submitted to re-extraction following the same procedure to generate supernatant B. Supernatant A and B were mixed and transferred to a clean 1.5 mL microtube with the appropriated volume of 15% NH_4_CO_3_. The samples were stored in a - 80° C freezer until analysis by LC-MS.

To extract polar metabolites from mouse plasma 40 µL of cold methanol was added to a 15 µL of mouse plasma in a 1.5 mL microtube. This mixture was vortexed for 10 seconds and incubated for 20 minutes in a −20° C freezer. Samples were centrifuged for 10 min at 16,000g at 4° C. Next, supernatant A was collected in a new tube, and the pellet was saved for re-extraction. For re-extraction, the pellet was resuspended in 200 µL of 40:40:20 buffer, vortexed, and allowed to sit on ice for 10 min. The samples were centrifuged for 10 min at 16,000 g at 4° C. Supernatant B was collected and mixed with supernatant A. This mixture was further processed to remove phospholipids using 1 mL Phenomenex (Phenomenex Inc.) tubes according to the manufacturer’s instruction and stored at −80° C until analysis by LC-MS.

The energy charge was calculated through the equation [ATP]+1/2[ADP] / [ATP]+[ADP]+[AMP] using the ion counts provided by metabolomics.

### Isotope tracing

KP and PGKP animals were put in an AA-crt or serine/glycine-free diet one week before NSCLC induction. After 11 weeks following NSCLC induction, the animals were subjected to aseptic surgical jugular vein catheterization (Instech, C20PV-MJV1301, 2 French, 10 cm) to implant the catheter to a vascular access button (Instech, VABM1B/25, 25 gauge, one-channel button) implanted under the back skin of the animal. The animals were left to recover for five days, according to the procedure previously described^3^. A fully recovered mouse was fasted for six hours and placed in a plastic harness (SAI Infusion Technologies), and the catheter was connected to an infusion pump (New Era Pump System) through a mouse tether and swivel system (Instech Laboratories). [U-^13^C]Glucose isotope tracer (110187-42.3 Cambridge Isotope Laboratories) was dissolved in sterile saline and infused at a rate of 3.5 nmol/g/min (0.1 μl/g/min) for 2.5h without a priming dose. Mice were euthanized after infusion and serum, tumor, lung, liver, and muscle (gastrocnemius) was collected. For each animal, one aliquot of serum, liver, and muscle was used to extract metabolites. For lung were used two aliquots from different lung lobes and tumors.

For the normalization of labeled isotopologue metabolites, ion counts per isotopologue were used to determine the percentage of each labeled isotopologue per metabolite 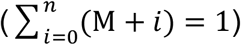. The obtained results were normalized by plasma glucose enrichment level determined by: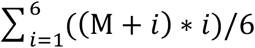.

### LC-MS/MS analyses

Metabolomics Shared Resource performed the LC-MS/MS on a Q Exactive PLUS hybrid quadrupole-orbitrap mass spectrometer coupled to a Vanquish Horizon UHPLC system (Thermo Fisher) with an XBridge BEH Amide column (150 mm × 2.1 mm, 2.5 μm particle size, Waters, Milford, MA). The HILIC separation used a gradient of solvent A (95%:5% H_2_O:acetonitrile with 20 mM acetic acid, 40 mM ammonium hydroxide, (pH 9.4) and solvent B (20%:80% H_2_O:acetonitrile with 20 mM acetic acid, 40 mM ammonium hydroxide, pH 9.4). The gradient was 0 min, 100% B; 3 min, 100% B; 3.2 min, 90% B; 6.2 min, 90% B; 6.5 min, 80% B; 10.5 min, 80% B; 10.7 min, 70% B; 13.5 min, 70% B; 13.7 min, 45% B; 16 min, 45% B; 16.5 min, 100% B; and 22 min, 100% B^96^. The flow rate was 300 μL/min. The column temperature was set to 25 °C. The autosampler temperature was set to 4 °C, and the injection volume was 5 μL. MS scans were obtained in negative ionization mode with a resolution of 70,000 at m/z 200 and an automatic gain control target of 3 x 106, and m/z scan range of 72 to 1000. The itaconate was monitored by a PRM event (m/z 129.02@HCD53.33). Metabolite data was obtained using the MAVEN software package^97^ (mass accuracy window: 5 ppm).

### Pathways impact analysis and metabolites classification

To evaluate the impact of the metabolic alterations found in KP versus PGKP TDCLs, we submitted the raw data obtained from the metabolic extraction on MAVEN (ion counts) to log transformation using the web-based software MetaboAnalyst 5.0.77. The transformed data were used to obtain the class of the metabolites significantly altered and the impact of these alterations on TDCLs^98^.

### Software, graphs, and figures

ImageJ 1.53J was used for image quantification using densitometry and labeled cell count. R 4.2.1 studio was used to generate heat maps. All charts and statistics presented in this article were built using GraphPad Prism 10 (GraphPad Prism, RRID: SCR_002798). The metabolites pick extraction was performed in El-Maven v0.12.0^99^. For tumor burden and quantification was used OLYMPUS OlyVIA 2.9. During the manuscript generation ChatGPT 3.4 (Open AI) was used as source of vocabulary, language, and readability. All Illustrations were drawn using Bio Render Premium and Adobe Illustrator 2023 (Adobe Illustrator, RRID:SCR_010279).

### Statistics and reproducibility

All statistical analyses were performed using GraphPad Prism 10.0 and MetaboAnalyst 5.0^98^. Samples sizes were chosen in advance based on the common practice of the described experiment and indicated by *n* or the number of points in the graph. Moreover, the initial sample size was determined based on a vast experience with tumor growth, survival, and manipulation of KP animals. The sample sizes for repeated experiments were based on the first one performed. Each experiment with mice was conducted with biological (at least three animals each) and technical (repeated at least twice) replicates. The experiments with TDCLs were also conducted with biological (at least three independent clones per genotype), technical (at least two independents well seeded per clone), and experimental (repeated at least twice) replicates. Based on Grubb’s test, outlier data were automatically excluded (GraphPad Prism). Outliers were removed from the plasma metabolic panel. All experiments randomly allocated mice into experimental groups. The investigators were not blinded during the experiments and outcome assessments. Blinding was not performed as the genotype and sex of the mice required identification for housing purposes. Sex differences were not identified among the genotypes and experiments.

## REFERENCES

1. Hanahan, D. & Weinberg, R. A. Hallmarks of cancer: The next generation. Cell vol. 144 646–674 Preprint at (2011).

2. Poillet-Perez, L. et al. Autophagy promotes growth of tumors with high mutational burden by inhibiting a T-cell immune response. Nature Cancer 2020 1:9 1, 923–934 (2020).

3. Poillet-Perez, L. et al. Autophagy maintains tumour growth through circulating arginine. Nature 2018 563:7732 563, 569–573 (2018).

4. Cararo-Lopes, E. et al. Integrated metabolic and genetic analysis reveals distinct features of human differentiated thyroid cancer. Clin Transl Med 13, e1298 (2023).

5. Tajan, M. et al. Serine synthesis pathway inhibition cooperates with dietary serine and glycine limitation for cancer therapy. Nature Communications 2021 12:1 12, 1–16 (2021).

6. Labuschagne, C. F., van den Broek, N. J. F., Mackay, G. M., Vousden, K. H. & Maddocks, O. D. K. Serine, but Not Glycine, Supports One-Carbon Metabolism and Proliferation of Cancer Cells. Cell Rep 7, 1248–1258 (2014).

7. Bartman, C. R. et al. Slow TCA flux and ATP production in primary solid tumours but not metastases. Nature 614, 349–357 (2023).

8. Bhatt, V. et al. Inhibition of autophagy and MEK promotes ferroptosis in Lkb1-deficient Kras-driven lung tumors. Cell Death & Disease 2023 14:1 14, 1–14 (2023).

9. Lan, T. et al. G6PD Maintains Redox Homeostasis and Biosynthesis in LKB1-Deficient KRAS-Driven Lung Cancer. bioRxiv (2023) doi:10.1101/2023.10.06.561131.

10. Liu, Y. & Birsoy, K. Metabolic sensing and control in mitochondria. Mol Cell 83, 877–889 (2023).

11. Zong, W. X., Rabinowitz, J. D. & White, E. Mitochondria and Cancer. Mol Cell 61, 667–676 (2016).

12. Liu, Q. et al. Metformin inhibits prostate cancer progression by targeting tumor-associated inflammatory infiltration. Clinical Cancer Research 24, 5622–5634 (2018).

13. Hirsch, H. A., Iliopoulos, D., Tsichlis, P. N. & Struhl, K. Metformin selectively targets cancer stem cells, and acts together with chemotherapy to block tumor growth and prolong remission. Cancer Res 69, 7507–7511 (2009).

14. Zarou, M. M., Vazquez, A. & Vignir Helgason, G. Folate metabolism: a re-emerging therapeutic target in haematological cancers. Leukemia 2021 35:6 35, 1539–1551 (2021).

15. Jordheim, L. P., Durantel, D., Zoulim, F. & Dumontet, C. Advances in the development of nucleoside and nucleotide analogues for cancer and viral diseases. Nature Reviews Drug Discovery 2013 12:6 12, 447–464 (2013).

16. Maddocks, O. D. K. et al. Modulating the therapeutic response of tumours to dietary serine and glycine starvation. Nature 2017 544:7650 544, 372–376 (2017).

17. Yang, M. & Vousden, K. H. Serine and one-carbon metabolism in cancer. Nature Reviews Cancer 2016 16:10 16, 650–662 (2016).

18. Ducker, G. S. & Rabinowitz, J. D. One-Carbon Metabolism in Health and Disease. Cell Metab 25, 27–42 (2017).

19. Amelio, I., Cutruzzolá, F., Antonov, A., Agostini, M. & Melino, G. Serine and glycine metabolism in cancer. Trends Biochem Sci 39, 191–198 (2014).

20. Lee, Y. S., Johnson, K. A., Molineux, I. J. & Yin, Y. W. A single mutation in human mitochondrial DNA polymerase Pol γA affects both polymerization and proofreading activities of only the holoenzyme. Journal of Biological Chemistry 285, 28105–28116 (2010).

21. Stumpf, J. D. & Copeland, W. C. Mitochondrial DNA replication and disease: Insights from DNA polymerase γ mutations. Cellular and Molecular Life Sciences 68, 219–233 (2011).

22. Weinberg, S. E. & Chandel, N. S. Targeting mitochondria metabolism for cancer therapy. Nat Chem Biol 11, 9 (2015).

23. Wallace, D. C. A Mitochondrial Paradigm of Metabolic and Degenerative Diseases, Aging, and Cancer: A Dawn for Evolutionary Medicine. 10.1146/annurev.genet.39.110304.095751 39, 359–407 (2005).

24. Kujoth, G. C. et al. Mitochondrial DNA Mutations, Oxidative Stress, and Apoptosis in Mammalian Aging. 309, 481–485 (2005).

25. Cancer Statistics Review, 1975-2017 - SEER Statistics. https://seer.cancer.gov/archive/csr/1975_2017/.

26. Prag, H. A. & Murphy, M. P. mtDNA mutations help support cancer cells. Nature Cancer 2020 1:10 1, 941–942 (2020).

27. Smith, A. L. M. et al. Age-associated mitochondrial DNA mutations cause metabolic remodeling that contributes to accelerated intestinal tumorigenesis. Nature Cancer 2020 1:10 1, 976–989 (2020).

28. Yuan, Y. et al. Comprehensive molecular characterization of mitochondrial genomes in human cancers. Nature Genetics 2020 52:3 52, 342–352 (2020).

29. Dai, Y. et al. Behavioral and metabolic characterization of heterozygous and homozygous POLG mutator mice. Mitochondrion 13, 282–291 (2013).

30. International Agency for Research on Cancer. IARC Biennial Report 2020-2021. IARC Publications (2021).

31. Travis, W. D. et al. The 2015 World Health Organization Classification of Lung Tumors: Impact of Genetic, Clinical and Radiologic Advances Since the 2004 Classification. J Thorac Oncol 10, 1243–1260 (2015).

32. DuPage, M., Dooley, A. L. & Jacks, T. Conditional mouse lung cancer models using adenoviral or lentiviral delivery of Cre recombinase. Nature Protocols 2009 4:7 4, 1064–1072 (2009).

33. George, J. et al. Comprehensive genomic profiles of small cell lung cancer. Nature 2015 524:7563 524, 47–53 (2015).

34. Trifunovic, A. et al. Premature ageing in mice expressing defective mitochondrial DNA polymerase. Nature 429, 417–423 (2004).

35. Kujoth, G. C. Mitochondrial DNA Mutations, Oxidative Stress, and Apoptosis in Mammalian Aging. 481, 481–484 (2007).

36. Li-Harms, X. J. et al. Mito-protective autophagy is impaired in erythroid cells of aged mtDNA-mutator mice. Blood 125, 162–174 (2015).

37. Chen, M. L. et al. Erythroid dysplasia, megaloblastic anemia, and impaired lymphopoiesis arising from mitochondrial dysfunction. 114, 4045–4054 (2017).

38. Vermulst, M. et al. Mitochondrial point mutations do not limit the natural lifespan of mice. Nat Genet 39, 540–543 (2007).

39. Goodman, R. P. et al. Hepatic NADH reductive stress underlies common variation in metabolic traits. Nature 583, 122–126 (2020).

40. Titov, D. V. et al. Complementation of mitochondrial electron transport chain by manipulation of the NAD+/NADH ratio. Science (1979) 352, 231–235 (2016).

41. Wu, J., Jin, Z., Zheng, H. & Yan, L. J. Sources and implications of NADH/NAD+ redox imbalance in diabetes and its complications. Diabetes, Metabolic Syndrome and Obesity 9, 145–153 (2016).

42. Xiao, W., Wang, R. S., Handy, D. E. & Loscalzo, J. NAD(H) and NADP(H) Redox Couples and Cellular Energy Metabolism. Antioxid Redox Signal 28, 251–272 (2018).

43. Mohr, S., Stamler, J. S. & Brüne, B. Posttranslational modification of glyceraldehyde-3-phosphate dehydrogenase by S-nitrosylation and subsequent NADH attachment. J Biol Chem 271, 4209–4214 (1996).

44. Tristan, C., Shahani, N., Sedlak, T. W. & Sawa, A. The diverse functions of GAPDH: Views from different subcellular compartments. Cell Signal 23, 317–323 (2011).

45. Zhu, X., Jin, C., Pan, Q. & Hu, X. Determining the quantitative relationship between glycolysis and GAPDH in cancer cells exhibiting the Warburg effect. Journal of Biological Chemistry 296, 100369–100370 (2021).

46. Locasale, J. W. et al. Phosphoglycerate dehydrogenase diverts glycolytic flux and contributes to oncogenesis. Nature Genetics 2011 43:9 43, 869–874 (2011).

47. Tajan, M. et al. Serine synthesis pathway inhibition cooperates with dietary serine and glycine limitation for cancer therapy. Nature Communications 2021 12:1 12, 1–16 (2021).

48. Hennequart, M. et al. The impact of physiological metabolite levels on serine uptake, synthesis and utilization in cancer cells. Nature Communications 2021 12:1 12, 1–10 (2021).

49. Zhou, X. et al. Serine alleviates oxidative stress via supporting glutathione synthesis and methionine cycle in mice. Mol Nutr Food Res 61, 1700262 (2017).

50. Alexandrov, L. B. et al. Signatures of mutational processes in human cancer. Nature 2013 500:7463 500, 415–421 (2013).

51. Sim, N. L. et al. SIFT web server: predicting effects of amino acid substitutions on proteins. Nucleic Acids Res 40, W452 (2012).

52. Lin, C. S. et al. Mouse mtDNA mutant model of Leber hereditary optic neuropathy. Proc Natl Acad Sci U S A 109, 20065–20070 (2012).

53. López-Otín, C., Blasco, M. A., Partridge, L., Serrano, M. & Kroemer, G. The Hallmarks of Aging. Cell 153, 1194–1217 (2013).

54. Guo, J. Y. et al. Autophagy provides metabolic substrates to maintain energy charge and nucleotide pools in Ras-driven lung cancer cells. Genes Dev 30, 1704–1717 (2016).

55. Karsli-Uzunbas, G. et al. Autophagy is required for glucose homeostasis and lung tumor maintenance. Cancer Discov 4, 915–927 (2014).

56. Lim, E. W. et al. Progressive alterations in amino acid and lipid metabolism correlate with peripheral neuropathy in Polg D257A mice. Sci. Adv 7, 4077–4092 (2021).

57. Saleem, A. et al. Polymerase gamma mutator mice rely on increased glycolytic flux for energy production. Mitochondrion 21, 19–26 (2015).

58. Birsoy, K. et al. An Essential Role of the Mitochondrial Electron Transport Chain in Cell Proliferation Is to Enable Aspartate Synthesis. Cell 162, 540–551 (2015).

59. Mattaini, K. R., Sullivan, M. R. & Vander Heiden, M. G. The importance of serine metabolism in cancer. Journal of Cell Biology 214, 249–257 (2016).

60. Bhatt, V. et al. Inhibition of autophagy and MEK promotes ferroptosis in Lkb1-deficient Kras-driven lung tumors. Cell Death & Disease 2023 14:1 14, 1–14 (2023).

61. Faubert, B., Tasdogan, A., Morrison, S. J., Mathews, T. P. & DeBerardinis, R. J. Stable isotope tracing to assess tumor metabolism in vivo. Nat Protoc 16, 5123–5145 (2021).

62. Diehl, F. F., Lewis, C. A., Fiske, B. P. & Vander Heiden, M. G. Cellular redox state constrains serine synthesis and nucleotide production to impact cell proliferation. Nature Metabolism 2019 1:9 1, 861–867 (2019).

63. Arnold, P. K. et al. A non-canonical tricarboxylic acid cycle underlies cellular identity. Nature 2022 603:7901 603, 477–481 (2022).

64. Murphy, J. P. et al. The NAD + Salvage Pathway Supports PHGDH-Driven Serine Biosynthesis. Cell Rep 24, 2381-2391.e5 (2018).

65. Pacold, M. E. et al. A PHGDH inhibitor reveals coordination of serine synthesis and one-carbon unit fate. Nature Chemical Biology 2016 12:6 12, 452–458 (2016).

66. Newman, A. C. & Maddocks, O. D. K. One-carbon metabolism in cancer. British Journal of Cancer 2017 116:12 116, 1499–1504 (2017).

67. Schober, F. A. et al. The one-carbon pool controls mitochondrial energy metabolism via complex i and iron-sulfur clusters. Sci Adv 7, (2021).

68. Ribas, V., García-Ruiz, C. & Fernández-Checa, J. C. Glutathione and mitochondria. Front Pharmacol 5 JUL, 151 (2014).

69. Weinberg, F. et al. Mitochondrial metabolism and ROS generation are essential for Kras-mediated tumorigenicity. Proc Natl Acad Sci U S A 107, 8788–93 (2010).

70. Cararo-Lopes, E. et al. Autophagy buffers Ras-induced genotoxic stress enabling malignant transformation in keratinocytes primed by human papillomavirus. Cell Death & Disease 2021 12:2 12, 1–16 (2021).

71. Han, S. H. et al. Mitochondrial integrated stress response controls lung epithelial cell fate. Nature 620, 890–897 (2023).

72. Poor, T. A. & Chandel, N. S. SnapShot: Mitochondrial signaling. Mol Cell 83, 1012-1012.e1 (2023).

73. Zong, W. X., Rabinowitz, J. D. & White, E. Mitochondria and Cancer. Mol Cell 61, 667–676 (2016).

74. Weinberg, F. et al. Mitochondrial metabolism and ROS generation are essential for Kras-mediated tumorigenicity. Proc Natl Acad Sci U S A 107, 8788–93 (2010).

75. Gopal, R. K. et al. Early loss of mitochondrial complex I and rewiring of glutathione metabolism in renal oncocytoma. Proc Natl Acad Sci U S A 115, E6283–E6290 (2018).

76. Joshi, S. et al. The Genomic Landscape of Renal Oncocytoma Identifies a Metabolic Barrier to Tumorigenesis Article The Genomic Landscape of Renal Oncocytoma Identifies a Metabolic Barrier to Tumorigenesis. CellReports 13, 1895–1908 (2015).

77. Zecchini, V. et al. Fumarate induces vesicular release of mtDNA to drive innate immunity. Nature 615, 499–506 (2023).

78. Tallini, G. Oncocytic tumours. Virchows Archiv 433, 5–12 (1998).

79. Stratton, M. R., Campbell, P. J. & Futreal, P. A. The cancer genome. Nature 2009 458:7239 458, 719–724 (2009).

80. Berridge, M. V., Dong, L. & Neuzil, J. Mitochondrial DNA in Tumor Initiation, Progression, and Metastasis: Role of Horizontal mtDNA Transfer. Cancer Res 75, 3203–3208 (2015).

81. Vasan, K., Werner, M. & Chandel, N. S. Mitochondrial Metabolism as a Target for Cancer Therapy. Cell Metab 32, 341–352 (2020).

82. Martínez-Reyes, I. et al. Mitochondrial ubiquinol oxidation is necessary for tumour growth. Nature 585, 288–292 (2020).

83. Gui, D. Y. et al. Environment Dictates Dependence on Mitochondrial Complex I for NAD+ and Aspartate Production and Determines Cancer Cell Sensitivity to Metformin. Cell Metab 24, 716–727 (2016).

84. Hosios, A. M. & Matthew, G. V. H. The redox requirements of proliferating mammalian cells. Journal of Biological Chemistry 293, 7490–7498 (2018).

85. Engel, A. L. et al. Serine-dependent redox homeostasis regulates glioblastoma cell survival. British Journal of Cancer 2020 122:9 122, 1391–1398 (2020).

86. Tajan, M. et al. Serine synthesis pathway inhibition cooperates with dietary serine and glycine limitation for cancer therapy. Nature Communications 2021 12:1 12, 1–16 (2021).

87. Reid, M. A. et al. Serine synthesis through PHGDH coordinates nucleotide levels by maintaining central carbon metabolism. Nature Communications 2018 9:1 9, 1–11 (2018).

88. Babraham Bioinformatics - FastQC A Quality Control tool for High Throughput Sequence Data. https://www.bioinformatics.babraham.ac.uk/projects/fastqc/.

89. Bolger, A. M., Lohse, M. & Usadel, B. Trimmomatic: a flexible trimmer for Illumina sequence data. Bioinformatics 30, 2114–2120 (2014).

90. Li, H. Aligning sequence reads, clone sequences and assembly contigs with BWA-MEM. (2013).

91. Okonechnikov, K., Conesa, A. & García-Alcalde, F. Qualimap 2: advanced multi-sample quality control for high-throughput sequencing data. Bioinformatics 32, 292–294 (2016).

92. ensemblorg/ensembl-vep - Docker Image | Docker Hub. https://hub.docker.com/r/ensemblorg/ensembl-vep.

93. Cingolani, P. et al. A program for annotating and predicting the effects of single nucleotide polymorphisms, SnpEff: SNPs in the genome of Drosophila melanogaster strain w1118; iso-2; iso-3. Fly (Austin) 6, 80 (2012).

94. Kuilman, T. et al. CopywriteR: DNA copy number detection from off-target sequence data. Genome Biol 16, 1–15 (2015).

95. Guo, J. Y. et al. Activated Ras requires autophagy to maintain oxidative metabolism and tumorigenesis. Genes Dev 25, 460–470 (2011).

96. Su, X. et al. In-Source CID Ramping and Covariant Ion Analysis of Hydrophilic Interaction Chromatography Metabolomics. Anal Chem 92, 4829–4837 (2020).

97. Melamud, E., Vastag, L. & Rabinowitz, J. D. Metabolomic analysis and visualization engine for LC - MS data. Anal Chem 82, 9818–9826 (2010).

98. Chong, J., Wishart, D. S. & Xia, J. Using MetaboAnalyst 4.0 for Comprehensive and Integrative Metabolomics Data Analysis. Curr Protoc Bioinformatics 68, e86 (2019).

99. Agrawal, S. et al. El-MAVEN: A Fast, Robust, and User-Friendly Mass Spectrometry Data Processing Engine for Metabolomics. Methods Mol Biol 1978, 301–321 (2019).

